# Differential Toxicity of Perfluorooctane Sulfonate (PFOS) in Wild-Type and Oatp1d1 Mutant Zebrafish Embryos

**DOI:** 10.1101/2024.06.28.601146

**Authors:** Ivan Mihaljevic, Lana Vujica, Jelena Dragojević, Jovica Lončar, Tvrtko Smital

## Abstract

This study presents a comprehensive analysis of the effects of perfluorooctane sulfonate (PFOS) exposure on zebrafish embryos, focusing on the differential responses between wild-type (WT) and oatp1d1 mutant embryos. The findings improve our understanding of the toxicokinetic and toxicodynamic mechanisms of PFOS, a persistent and bioaccumulative member of the per- and polyfluoroalkyl substances (PFAS) family. The study revealed significant differences in mortality rates with calculated LC50 values of 23.57 µM for WT and 16.71 µM for oatp1d1 mutants, indicating a higher susceptibility of the mutants to PFOS toxicity. This indicates the crucial role of the Oatp1d1 transporter in mediating the toxic effects of PFOS, possibly related to detoxification processes or regulation of bioavailability. Developmental abnormalities, particularly in the swim bladder, were more pronounced in mutant embryos, indicating the role of the transporter in mitigating PFOS-induced developmental toxicity. Gene expression analysis showed differential modulation of biotransformation genes, including cytochrome P450 (cyp) and glutathione S-transferase (gst) genes, underscoring the complexity of PFOS toxicity. The study also highlighted disruptions in lipid metabolism with altered expression of genes involved in lipid synthesis and oxidation, leading to abnormal lipid accumulation. These findings have significant implications for aquatic ecosystems and human health, as PFOS is persistent and bioaccumulative in the environment. The study emphasizes the need for stringent regulatory measures and effective remediation strategies to address PFOS contamination and protect both aquatic life and human populations.

## INTRODUCTION

Per- and polyfluoroalkyl substances (PFAS) are a diverse group of man-made chemicals that have been widely used in various industrial applications and consumer products since the 1940s. These substances are characterized by their strong carbon-fluorine bonds, which give them remarkable resistance to water, oil and heat. This resistance has made PFAS invaluable in numerous areas, such as non-stick cookware, water-repellent clothing, stain-resistant fabrics and carpets, food packaging, firefighting foams, and personal care products such as shampoos and dental floss (Buck et al., 2011). However, the same chemical stability that makes PFAS so useful also leads to environmental persistence and widespread contamination, raising significant ecological and health concerns (Giesy & Kannan, 2001Lau et al., 2007).

PFAS are not easily degradable and can persist in the environment for long periods of time (Prevedouros et al., 2006). They have been detected in various environmental media, including surface waters (rivers, lakes and oceans), groundwater and drinking water supplies, soils and sediments and even atmospheric dust (Ahrens, 2011; Post et al., 2012). Environmental concentrations of PFAS can vary widely, typically ranging from parts per trillion (ppt) to parts per billion (ppb), depending on proximity to sources such as industrial sites, firefighting training areas, and wastewater treatment plants (Wang et al., 2022; Sun et al., 2016). The persistence and bioaccumulative nature of PFAS results in their accumulation in the tissues of aquatic organisms, leading to higher concentrations in predators at the top of the food chain. This bioaccumulation poses a risk not only to wildlife, but also to humans who consume contaminated food (Consoer et al., 2016). PFAS exposure can result in various toxic effects on aquatic organisms, including developmental and reproductive toxicity, endocrine disruption, immune system effects and behavioral changes (Ankley et al., 2021; Houde et al., 2006). These toxic effects can lead to reduced hatching rates, abnormal development, reproductive failures, altered growth, metabolism and increased susceptibility to diseases. Additionally, PFAS exposure can result in behavioral changes, such as altered feeding and avoidance of predators, which can impact survival and reproduction (Lou et al., 2013).

Perfluorooctane sulfonate (PFOS) is a specific type of PFAS that has been widely used due to its hydrophobic and lipophobic properties. PFOS has been widely produced and utilized in various industrial and consumer products, including stain and water repellents for textiles, upholstery, carpets and leather products; firefighting foams used in aqueous film-forming foams (AFFF) for fire suppression, particularly at airports and military bases; coating and etching processes in the metal plating industry; electronics manufacturing processes such as photolithography; pesticides and insecticides; and paper and packaging materials to resist grease and moisture (Sato et al., 2009).

PFOS poses significant risks to aquatic organisms due to its persistence, bioaccumulative nature, and toxicity. PFOS readily bioaccumulates in aquatic organisms, leading to higher concentrations in top predators (Conder et al., 2008). Exposure to PFOS has been associated with various adverse effects on aquatic organisms, including developmental and reproductive toxicity, endocrine disruption, liver toxicity, immune system effects and behavioral changes (Ankley et al., 2021). These toxic effects can result in developmental abnormalities, reduced hatching success, impaired reproductive function, liver damage, altered lipid metabolism, increased susceptibility to infections and diseases, and changes in food intake, predator avoidance and mating behavior (Houde et al., 2006). The effects of PFOS on lipid metabolism, particularly in zebrafish embryos, involve several pathways and cellular processes. PFOS can interact with peroxisome proliferator-activated receptors (PPARs), particularly PPARα, resulting in altered expression of genes involved in fatty acid oxidation, lipid transport and lipid synthesis (Rosenmai et al., 2016). PFOS exposure induces oxidative stress through the generation of reactive oxygen species (ROS), which can damage cellular components, including lipids, leading to lipid peroxidation (Lau et al., 2007). PFOS-induced mitochondrial dysfunction can impair lipid metabolism, resulting in energy deficits and lipid accumulation (Domingo & Nadal, 2019). Additionally, PFOS can induce endoplasmic reticulum (ER) stress and impair protein folding and lipid homeostasis (Ankley et al., 2021). PFOS can disrupt the normal function of lipoproteins and enzymes involved in lipid transport and storage, leading to abnormal lipid accumulation or depletion in tissues (Houde et al., 2011). Furthermore, PFOS exposure can lead to changes in the expression of key genes involved in lipid metabolism and hormonal disruption, affecting lipid metabolic processes (Rosenmai et al., 2016).

Studies have shown that PFOS exposure leads to increased lipid accumulation in zebrafish embryos due to impaired lipid catabolism and enhanced lipid synthesis (Domingo & Nadal, 2019). PFOS exposure can change the composition of lipids in zebrafish embryos, including increased triglyceride and cholesterol levels (Lau et al., 2007). PFOS has been shown to upregulate genes involved in lipid synthesis and downregulate genes involved in lipid oxidation (Ankley et al., 2021).

PFAS, and particularly PFOS, represent a significant environmental and public health challenge due to their persistence, bioaccumulation and toxicity. The widespread use of PFAS in various applications has led to global contamination that requires continuous efforts to monitor and mitigate their impact. Understanding the mechanisms by which PFAS affects lipid metabolism and other biological processes is crucial for developing strategies to protect aquatic ecosystems and human health. Regulatory measures to limit PFAS use and to remediate contaminated sites are essential steps in addressing this pervasive environmental issue (Cousins et al., 2016).

PFOS interacts with various organic anion transporters (OATs) and organic anion transporting polypeptides (OATPs). These transporters play a crucial role in the uptake, distribution and excretion of PFOS in humans and animals. Research has shown that PFOS can be transported by several OATs, including hOAT4 (Nakagawa et al., 2009) and OATPs such as OATP1A2, OATP1B1 and OATP1B3, which are expressed in different tissues including the liver, kidney and intestine. Studies have shown that hOAT4 is involved in the renal clearance of PFOS and facilitates its excretion from the body (Nakagawa et al., 2009) Similarly, OATPs in the liver and intestine contribute to the disposition of PFOS by mediating its uptake into hepatocytes and enterocytes (Kimura et al., 2020). This transport activity affects the bioavailability and toxicity of PFOS, influencing its accumulation in tissues and its potential health effects (Zhao et al., 2016). An important aspect of PFOS interaction with these transporters is its impact on reproductive toxicity, particularly through OATP3a1 in Sertoli cells, which may lead to adverse effects on male reproductive health (Li et al., 2023). Additionally, the involvement of transporters such as NTCP and ASBT in the liver highlights the complexity of PFOS disposition and its potential impairment of bile acid transport (Zhao et al., 2015).

The toxicokinetics of PFOS in zebrafish are influenced by its interaction with various transporters, including Oatps and Oats. These interactions can lead to significant changes in the bioconcentration and tissue distribution of PFOS in zebrafish (Consoer et al., 2016). Zebrafish Oats, including Oat1 and Oat3, play an important role in modulating the transport activity of PFOS. Specifically, zebrafish Oat1 and Oat3 exhibit interactions with environmental contaminants, including PFOS, affecting their bioavailability and toxicity in zebrafish (Dragojević et al., 2020). Oatp1d1 has a high affinity towards PFOS. This interaction highlights the role of OATP1d1 in mediating the uptake and potential toxic effects of PFOS in zebrafish (Popovic et al., 2014). Additionally, the functional conservation of OATPs/Oatps in vertebrates suggests that zebrafish Oatp1d1 might serve as a functional ortholog to human OATP2B1, showing strong interactions with PFOS and impacting its toxicity (Dragojević et al., 2021). The expression of downstream pathways, including those involving Oatps, was significantly altered in zebrafish embryos, indicating the crucial role of these transporters in mediating the effects of PFOS (Jantzen et al., 2016).

In this study, we explored the impact of perfluorooctane sulfonate (PFOS) exposure on both wild-type (WT) and *oatp1d1* mutant embryos, focusing on mortality rates, developmental abnormalities, oxidative stress, apoptosis, and gene expression changes. Our results demonstrated that PFOS exposure led to disruptions in normal embryonic development, with distinct differences observed between WT and mutant embryos. Specifically, mutant embryos exhibited higher mortality rates, increased deformities in the swim bladder, and greater incidences of scoliosis and necrosis. Furthermore, elevated oxidative stress and apoptosis levels were more pronounced in mutant embryos, indicating a heightened sensitivity to PFOS. Gene expression analysis revealed differential regulation of several genes involved in biotransformation processes, with notable variations in *oatp1d1* and other related genes under various PFOS concentrations and exposure durations. These findings offer valuable insights into the toxicological effects of PFOS and the role of Oatp1d1 in the associated defense mechanisms.

## MATERIALS AND METHODS

### Ethics

This study was approved by the Ethics Committee of the Ruđer Bošković Institute, the National Committee for Animal Welfare, and the Ministry of Agriculture in Zagreb, Croatia (Permission no. 525-10/124120-9), and carried out in accordance with the European Communities Council Directive of September 22, 2010 (2010/63/EU).

### Chemicals

All tested compounds, model fluorescent substrates and interactors alike were purchased from Sigma-Aldrich (Taufkirchen, Germany).

### Animals and sample collection

Wild-type (WT) zebrafish, ABO strain (European Zebrafish Resource Centre, Karlsruhe, Germany), were maintained under standard conditions, with a 14-hour light/10-hour dark cycle and a water temperature of 27–28 °C. The fish were fed with a standard food of the appropriate size (Gemma Micro, Skretting, Norway). Embryos were obtained by morning spawning from 1 to 1.5 year old zebrafish, transferred to a petri dish containing E3 medium (5 mM NaCl, 0.17 mM KCl, 0.33 mM CaCl2, 0.33 mM MgSO4) and reared in an incubator at 28 °C with the same light/dark period as the adults. Embryonic development was observed under a dissecting microscope (Motic AE31E, Motic, Barcelona, Spain) and embryos were staged as previously described (Kimmel et al., 1995).

### Generation of the *oatp1d1* mutant zebrafish line

Oatp1d1 functional knockout zebrafish line was developed previously (Vujica et al., n.d.). In brief, a short guide (sg)RNA (5′GGACTCGCATTTGTAAGGCA3′) targeting exon four was selected using the CRISPR scan algorithm (Moreno-Mateos et al., 2015) and generated as previously described (Modzelewski et al., 2018) (Fig. 1A). A mixture of 600 ng/µL of Cas9 protein (NEB: M0386) and sgRNA complex was prepared at a 1:1 ratio and 1 nL of the mixture was injected into zebrafish embryos at the one-cell stage. When the injected fish (F0) reached adulthood at about three months of age, they were crossed with WT fish and their progeny were analyzed by high-resolution melting analysis (HRMA). Embryos positive for mutations were subsequently sequenced and several mutations were detected in the germline of the F0 fish. Female and male F0 fish that carried the same mutation (deletion of 5 nucleotides (GTGCC) at position 9115-9119) on the genomic DNA in the germline, resulting in a premature stop codon mutation at 140 amino acids, were selected as founders (Fig. S1). The founders were then crossed with the aim of producing homozygous mutant embryos. As the offspring grew, their fins were clipped to isolate the genetic material of each fish for genotyping (IACUC standard procedure, 2022). Female and male fish with homozygous mutation (premature stop codon at 140 amino acids) were selected for further crosses to generate F2 generations of fish lacking the functional Oatp1d1 protein. Genotyping by sequencing confirmed that all F2 fish were homozygous mutants with 1 or 2 mutations (Table S1).

**Figure 1.**
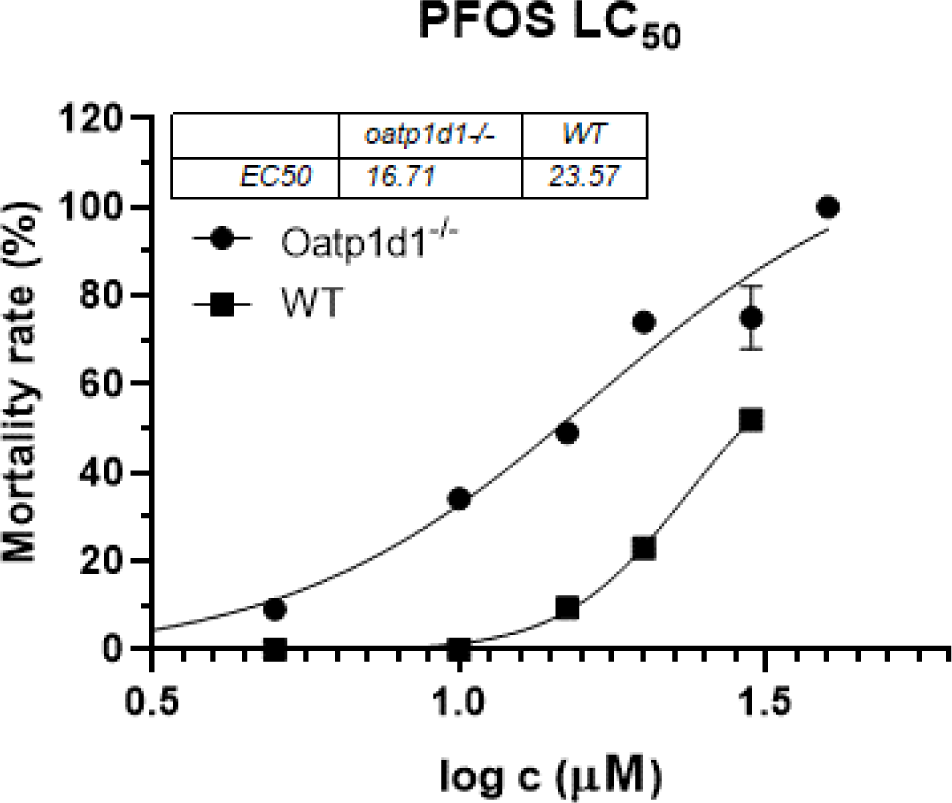
Dose–response (LC_50_) curves used for the calculations of the zebrafish WT and *oatp1d1* mutant embryos mortality after 96 hours of exposure to increasing PFOS concentrations. Data are results of a typical experiment performed in triplicate. Error bars indicate standard deviations (SD). Dose–response curves were generated using GraphPad Prism 8 software.

### PFOS exposure experiments with zebrafish embryos

To investigate the effects of PFOS (CAS: 1763-23-1) on embryonic development, we performed exposure experiments with PFOS concentrations ranging from 5 to 30 µM. PFOS stock solutions (100 mM) were prepared in dimethyl sulphoxide (DMSO) and working solutions were prepared in E3 medium (DMSO <0.05%). Negative controls were prepared with E3 medium and DMSO. Sets of ten embryos were distributed in 24-well plates containing 1 mL of E3 medium per well. Exposure to PFOS began on day 1 post-fertilization (dpf) or 24 hours post-fertilization (hpf) and continued for up to five dpf or three to four days post-exposure (dpe). During this time, mortality and developmental abnormalities caused by PFOS were recorded.

### Gene expression analysis

Tissues of at least three individual fish were stored in RNA later at -20 °C for RNA isolation. Zebrafish were anesthetized and sacrificed by prolonged immersion in ice/cold water. Embryos (three independent pools, 10 – 20 embryos per pool) were collected at different developmental stages (1 – 5 dpf), dry frozen and stored at -80 °C until RNA extraction. Tissues were homogenized with an Ultra Turrax T25 homogenizer at medium intensity for 10 s (sometimes two cycles of 10 s per sample, depending on the tissue) and embryos were homogenized with a pestle homogenizer. RNA was isolated using the Trizol method (Tri reagent), quality checked by gel electrophoresis (two bands) and quantified using the BioSpec nano micro-volume spectrophotometer (Shimadzu, Kyoto, Japan). Reverse transcription (1 µg total RNA or 500 ng at lower concentrations) was performed using the ProtoScript II First Strand cDNA Synthesis Kit (NEB, E6560L), resulting in cDNA concentrations of 50 and 25 ng/µL, respectively, and the High-Capacity cDNA Reverse Transcription Kit with RNase Inhibitor (Applied Biosystems, Foster City, CA, USA), which resulted in a cDNA concentration of 100 ng/µL. qPCR primers were previously described in (Mihaljevic et al., 2016; Popovic et al., 2013) and qPCR was performed using the GoTAQ qPCR mix (Promega, A6001). The elongation factor (*EF1α*) was chosen as the housekeeping gene for the tissues because its expression is similar in all tissues analyzed, and the *ATP50* gene was chosen as the housekeeping gene for the embryos for the same reasons. The qPCR reaction mixture was prepared to a final volume of 10 µL with 10 ng of cDNA per reaction, and relative quantification was performed as described previously (Lončar et al., 2010) except that data were presented as mean normalized expression multiplied by a factor of 106. Arbitrary thresholds for expression of transcripts in qRT-PCR data were defined as follows: Genes were considered minimally expressed if the MNE was <999 ×107, low if the MNE was 1000 - 4,999, mild if the MNE was 5,000 - 8,999, moderate if the MNE was 9000 - 19999, moderately high if the MNE was 20000 - 29999, high if the MNE was 30000 - 49999, and very high if it was above 50000.

### Acridine orange staining (negdje ga zovu i cell death assay)

Apoptotic cells in zebrafish embryos (apoptosis of zebrafish embryos?) were visualised using acridine orange (AO), a nucleic acid selective metachromatic dye that emits green fluorescence when intercalated with DNA. It is often used to detect apoptosis in zebrafish because it can permeate apoptotic cells and bind to DNA, whereas normal cells are nonpermeable to acridine orange (Asharani et al. 2008). After exposure to 15 µM PFOS, at 120 hpf (5 dpf) zebrafish embryos were selected and stained with AO (5 µg/mL) for 20 min in the dark according to the protocol (Hema et al 2023). Stained embryos were washed three times for five minutes in E3, anesthetised in 0.02 % Tricaine (MS222) and dead/apoptotic cells were visualized under a fluorescence microscope using a green fluorescence filter (excitation – 480 nm, emission – 535 nm).

### ROS assay

PFOS induced ROS formation was visualised using cell-permeant 2′,7′-dichlorodihydrofluorescein diacetate (H_2_DCFDA). Zebrafish embryos exposed to 5 µM PFOS after 120 hpf were randomly selected, washed with PBS and incubated in 10 µM H_2_DCFDA for 20 min in the dark. After the incubation, embryos were washed with PBS to remove the excess dye, anesthetised in 0.02 % Tricaine (MS-222) and observed under the fluorescence microscope using green fluorescence filter (excitation – 480 nm, emission – 535 nm).

### Larval behavior

Before the experiment, 1 dpf larvae were exposed to 10 µM PFOS. After 96 hours of exposure, a locomotor behavior test was conducted on both PFOS-treated and untreated groups and on both WT and *oatp1d1* mutant embryos. Initially, eight zebrafish larvae per condition were randomly selected and placed into a 96-well plate, with each well containing 150 µL of E3 culture medium. The larvae were then given 10 minutes to adapt to the light stimuli and the observation chamber. Subsequently, they were exposed to alternating light and dark cycles (5 minutes light, 5 minutes dark, repeated once more). The locomotor behaviors were recorded using the DanioVision observation chamber (Noldus Information and Technology, Wageningen, the Netherlands) for automated tracking. The data on locomotor activities were analyzed using EthoVision XT 10.0 software.

### Data analysis

The data were plotted and statistically analyzed using the unpaired two-sided Student’s t-test with GraphPad Prism 8. Significance was attributed to differences between two independent variables when the p-value was less than 0.05. All experiments were performed in two to five biological replicates and the means ±standard deviations (SD) or standard errors (SE). Fluorescence intensity of ROS generation was quantified by ImageJ software (ImageJ bundled with Java 1.8.0_172).

## RESULTS

### Exposure to PFOS

Exposure of WT and mutant embryos to PFOS resulted with changes in normal embryonic development. Increasing concentrations of PFOS resulted in lethal effects at later stages and higher concentrations, with some characteristic sublethal effects at lower concentration ranges. Mortality rates (LC_50_) differed between WT and mutant embryos with a calculated LC_50_ of 23.57 for WT and 16.71 µM for *oatp1d1* mutants (Fig. 1). The mortality rates of the unexposed and the DMSO-exposed negative control groups were below 5 % (Fig. S8).

The most prominent effects of PFOS exposure were visible from 4 dpf and later. PFOS affected normal development of the swim bladder, resulting in a nonexistent or very small swim bladder, which caused the embryos to be on their sides and be less active (Fig. 2). The percentage of embryos with deformed swim bladders was higher in *oatp1d1* mutant embryos and increased with higher concentrations of PFOS. This effect also contributed to the lower activities of 10 µM PFOS exposed mutant embryos. Additionally, the exposed mutant embryos developed necrosis in the head and intestinal regions, with small or absent swim bladders.

**Figure 2.**
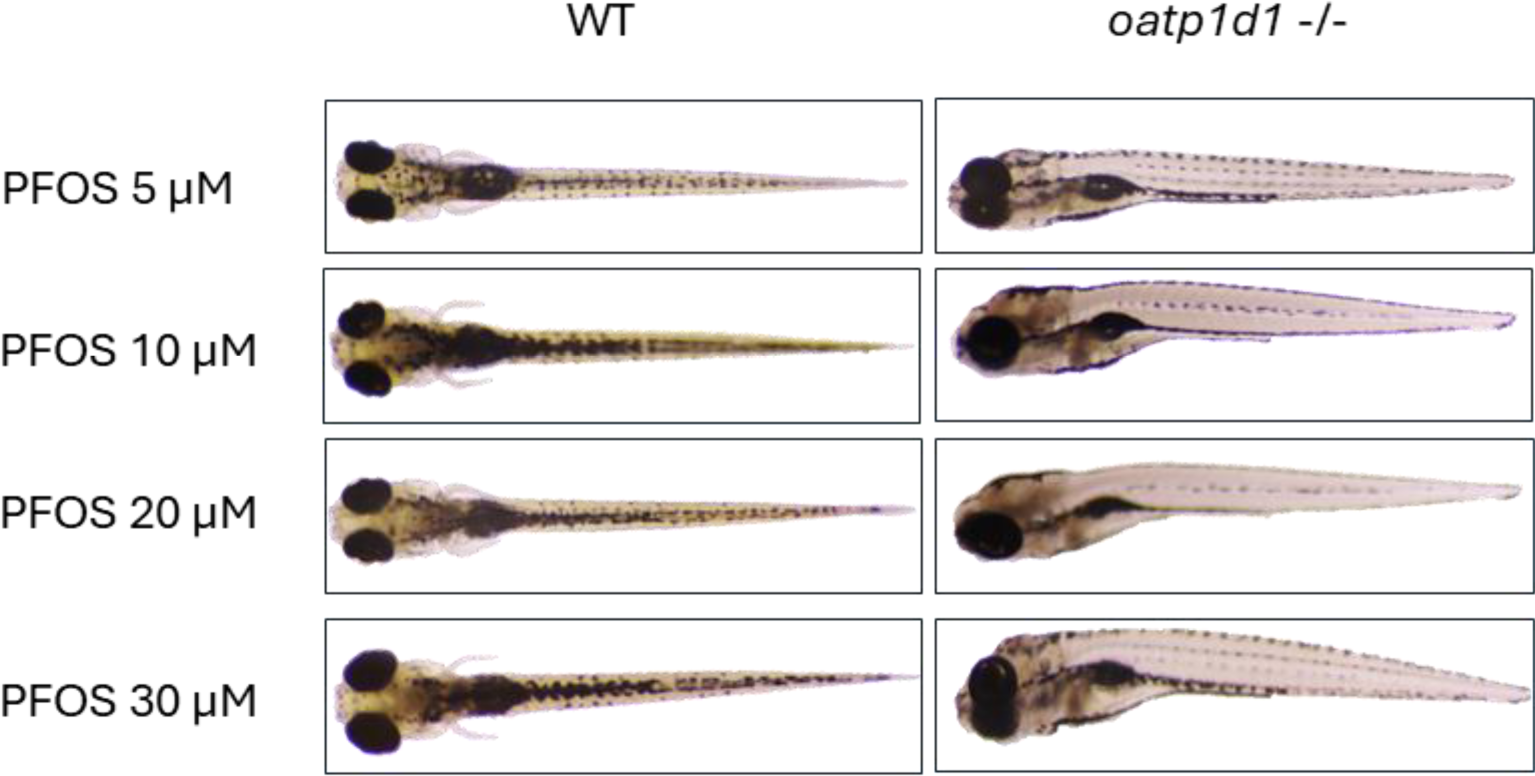
Effects of PFOS exposure on the development of *oatp1d1* mutant embryos (4 dpf). Onset of necrosis in the head and gut along with deformations of the swim bladder, resulting with mutant embryos being on their sides and less active.

At later stages, between 4 and 5 dpf, PFOS exposure caused more deleterious effects on embryonal development in the form of scoliosis, which was more present in the mutant embryos at the current developmental stages (4 - 5 dpf) (Fig. 3).

**Figure 3.**
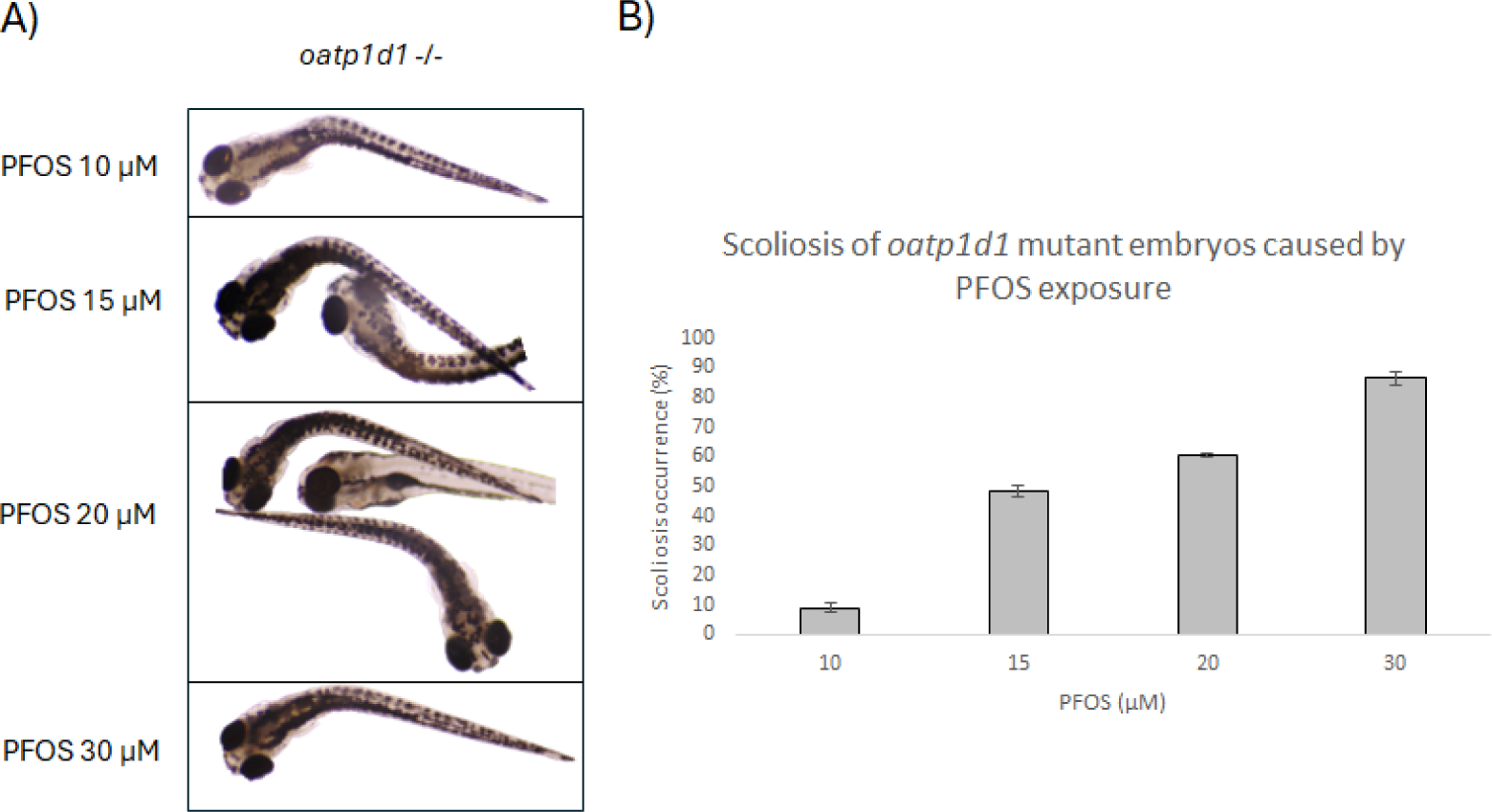
The occurrence of scoliosis in oatp1d1 mutant embryos as an effect of PFOS treatment. (A) Pictures showing the scoliosis effect in mutant embryos at 4 – 5 dpf. (B) Dose response of scoliosis occurrence in PFOS treated oatp1d1 mutant embryos.

Furthermore, PFOS exposure caused higher levels of oxidative stress, together with higher levels of apoptosis in *oatp1d1* mutant embryos. Exposure to 15 µM PFOS caused 2-fold higher levels of ROS species in *oatp1d1* mutant embryos at 5 dpf, whereas exposed WT and DMSO-exposed control embryos showed lower levels of oxidative stress. (Figs. 9 and S7). PFOS-exposed mutant embryos showed 3-fold higher levels of apoptosis compaing with WT and control embryos (Figs. 10 and S7).

PFOS exposure caused changes in the transcript expression of numerous genes involved in biotransformation processes. Oatp1d1 showed upregulation of 1.5-fold in 5 dpf wt embryos after exposure to 15 µM PFOS for 1 h, while the mutant embryos did not show any change (Fig. 5). Longer exposure to 5 µM PFOS did not cause any changes in the transcript expression of *oatp1d1* in both wt and mutant embryos (Fig. 6). However, at 4 dpf, wt embryos showed high downregulation of 0.19-fold upon exposure to 30 µM PFOS for three days (Fig. 7). On the other hand, a lower PFOS concentration (15 µM) caused a 2.16- and 3.23-fold upregulation of *oatp2b1* at 4 dpf and 3 days of exposure in both wt and mutant embryos, respectively (Fig. 8).

**Figure 4.**
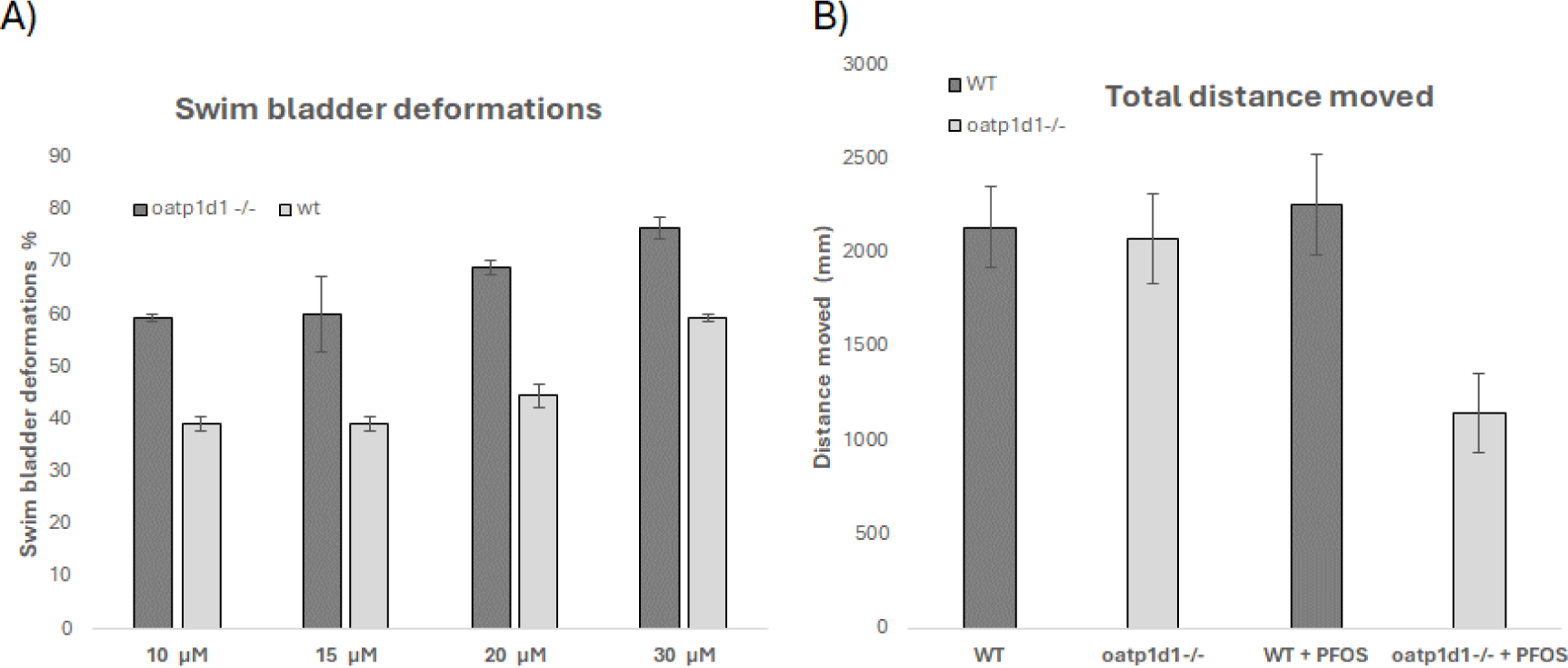
Effects of PFOS treatment on development of swim bladder in zebrafish embryos. (A) Dose dependent occurrence of swim bladder malformation in WT and mutant embryos. (B) Effects of PFOS treatment on movement of WT and mutant embryos.

**Figure 5.**
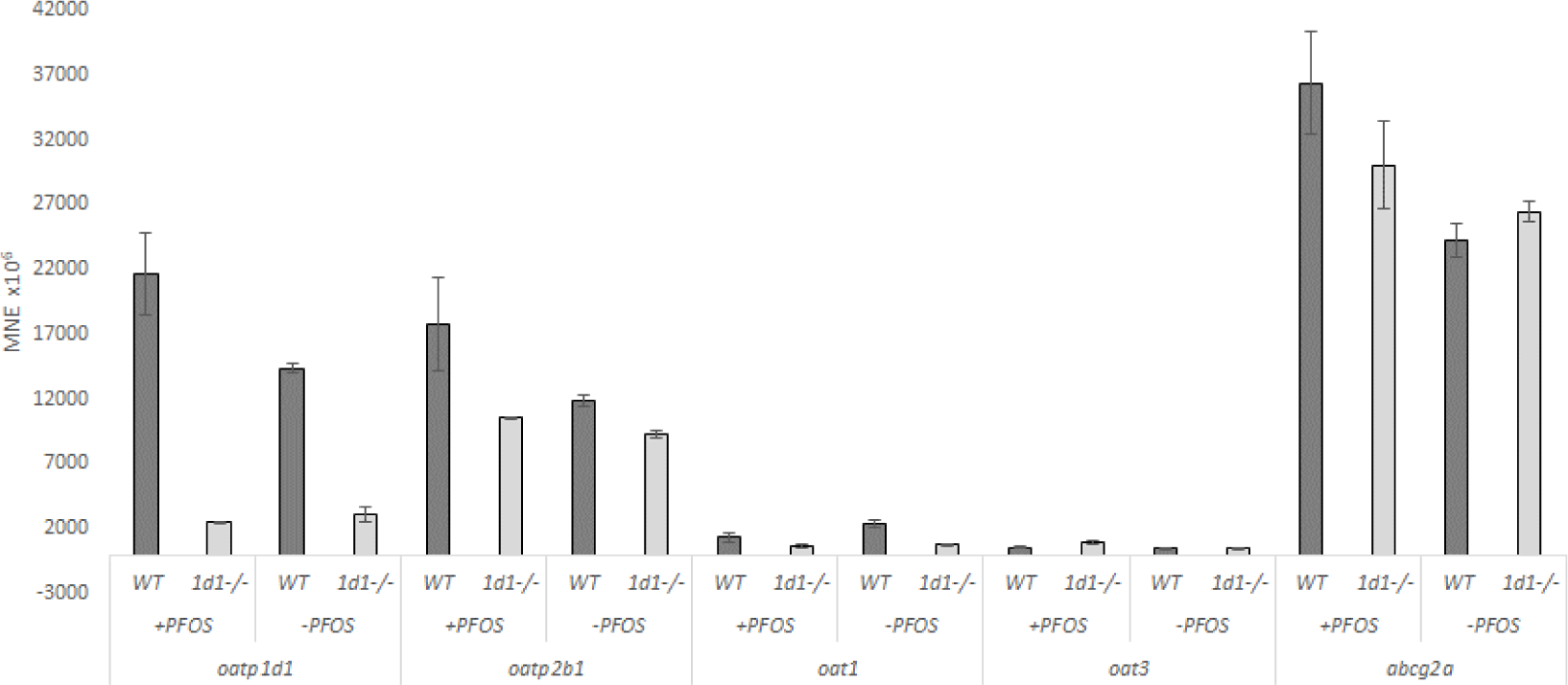
Transcript expression patterns of *oatp1d1, oatp2b1, oat1, oat3* and *abcg2a* in WT and *oatp1d1* mutant embryos at 5 dpf and with and without 1h exposure to 15 µM PFOS. Results of pools of 10 embryos per sample are presented. Data represents MNE (mean normalized expression) ± SD normalized to the housekeeping gene *ATP50*.

**Figure 6.**
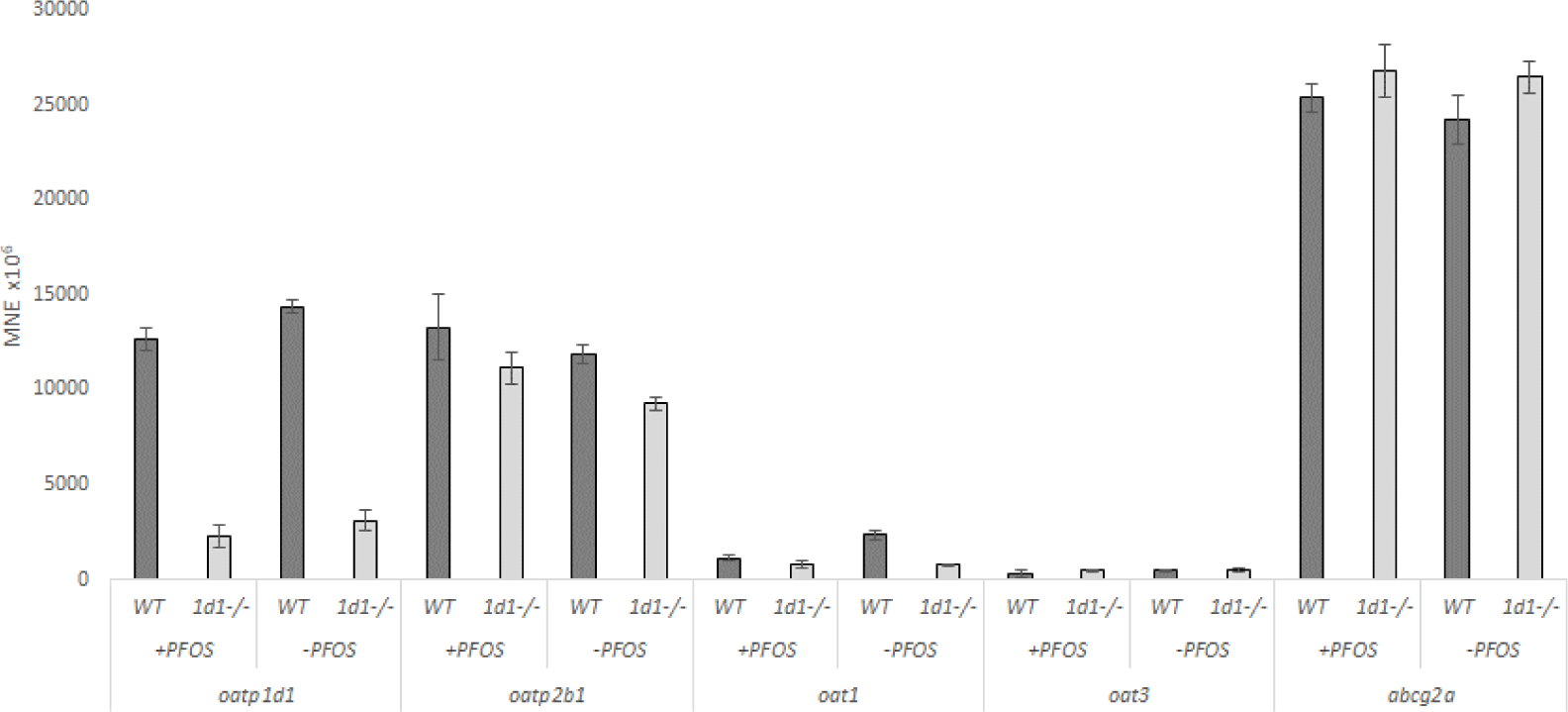
Transcript expression patterns of *oatp1d1, oatp2b1, oat1, oat3* and *abcg2a* in WT and *oatp1d1* mutant embryos at 5 dpf and with and without 4 days exposure to 5 µM PFOS. Results of pools of 10 embryos per sample are presented. Data represents MNE (mean normalized expression) ± SD normalized to the housekeeping gene *ATP50*.

**Figure 7.**
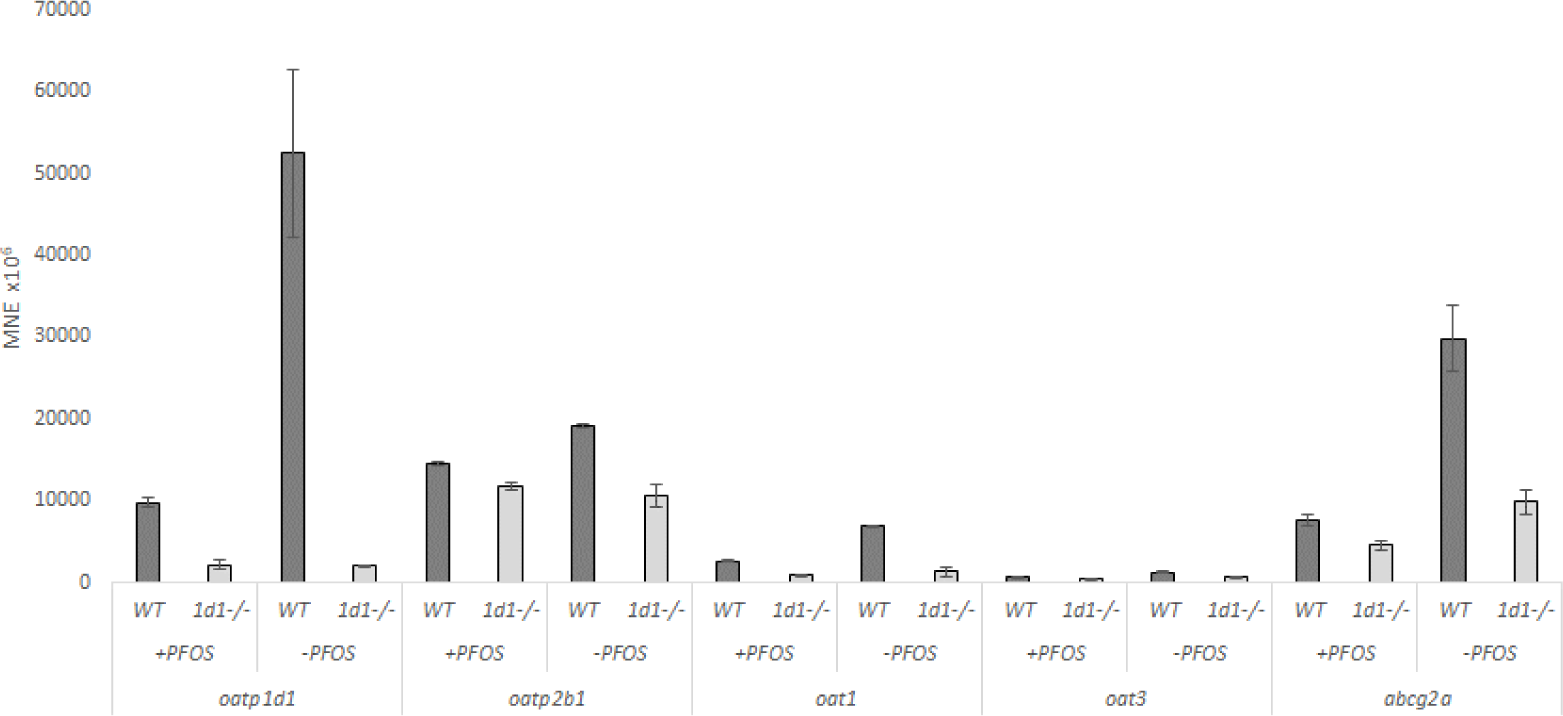
Transcript expression patterns of *oatp1d1, oatp2b1, oat1, oat3* and *abcg2a* in WT and *oatp1d1* mutant embryos at 4 dpf and with and without 3 days exposure to 30 µM PFOS. Results of pools of 10 embryos per sample are presented. Data represents MNE (mean normalized expression) ± SD normalized to the housekeeping gene *ATP50*.

**Figure 8.**
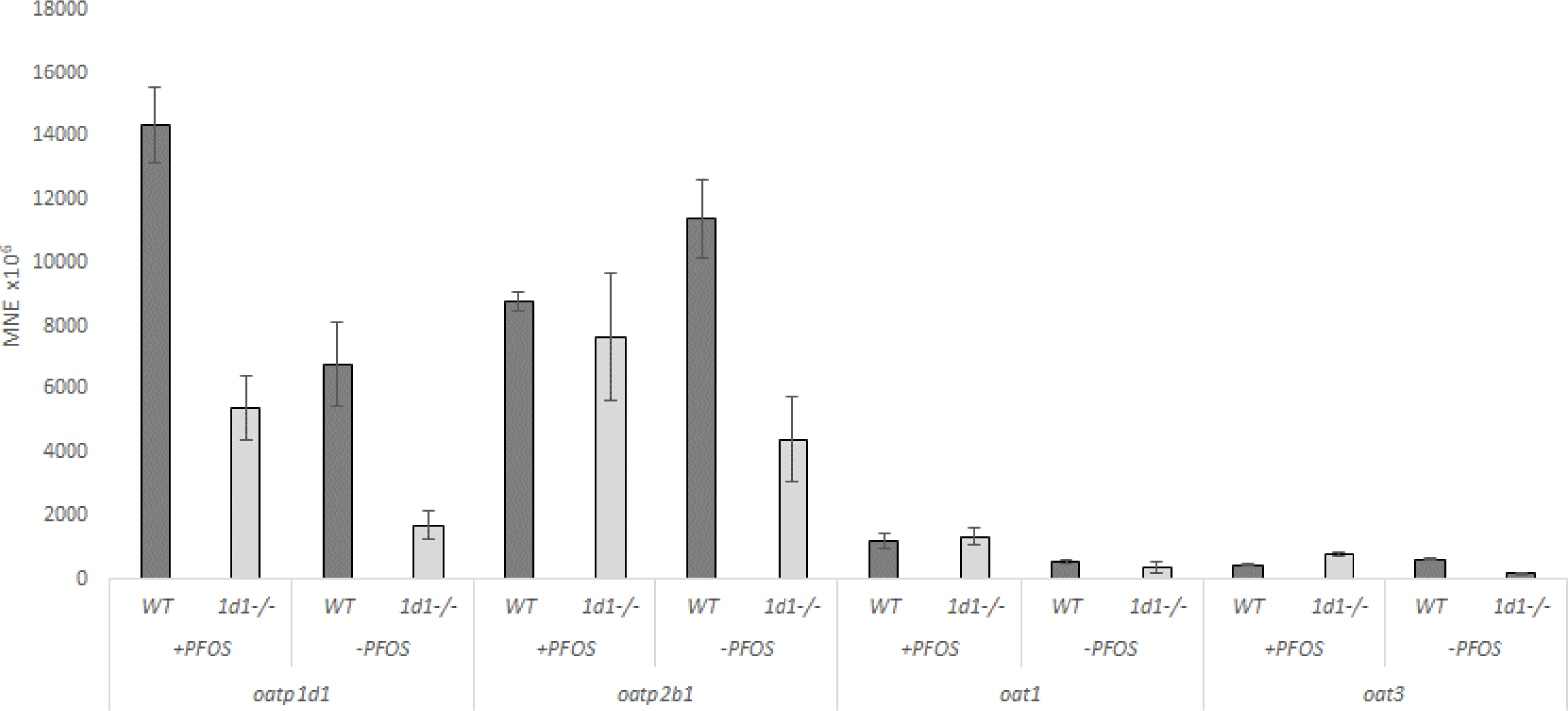
Transcript expression patterns of *oatp1d1, oatp2b1, oat1, oat3* and *abcg2a* in WT and *oatp1d1* mutant embryos at 4 dpf and with and without 3 days exposure to 15 µM PFOS. Results of pools of 10 embryos per sample are presented. Data represents MNE (mean normalized expression) ± SD normalized to the housekeeping gene *ATP50*.

*Oapt2b1* showed a 1.5-fold upregulation in 5 dpf wt embryos after exposure to 15 µM PFOS for 1 h, and a 1.14-fold upregulation in mutant embryos (Fig. 5). Longer exposure to 5 µM PFOS caused minor changes in oatp2b1 transcript expression in both wt and mutant embryos by 1.12- and 1.2-fold, respectively (Fig. 6). Wild type 4 dpf embryos after 3 days of exposure with 15 µM PFOS showed downregulation of 0.77-fold, whereas mutant embryos showed upregulation of 1.73-fold (Fig. 8).

Both oat1 and oat3 genes showed generally low expression under all conditions, with weak modulations of transcript expression upon exposure to different PFOS concentrations (Figs. 5, 7, 9, 10).

**Figure 9.**
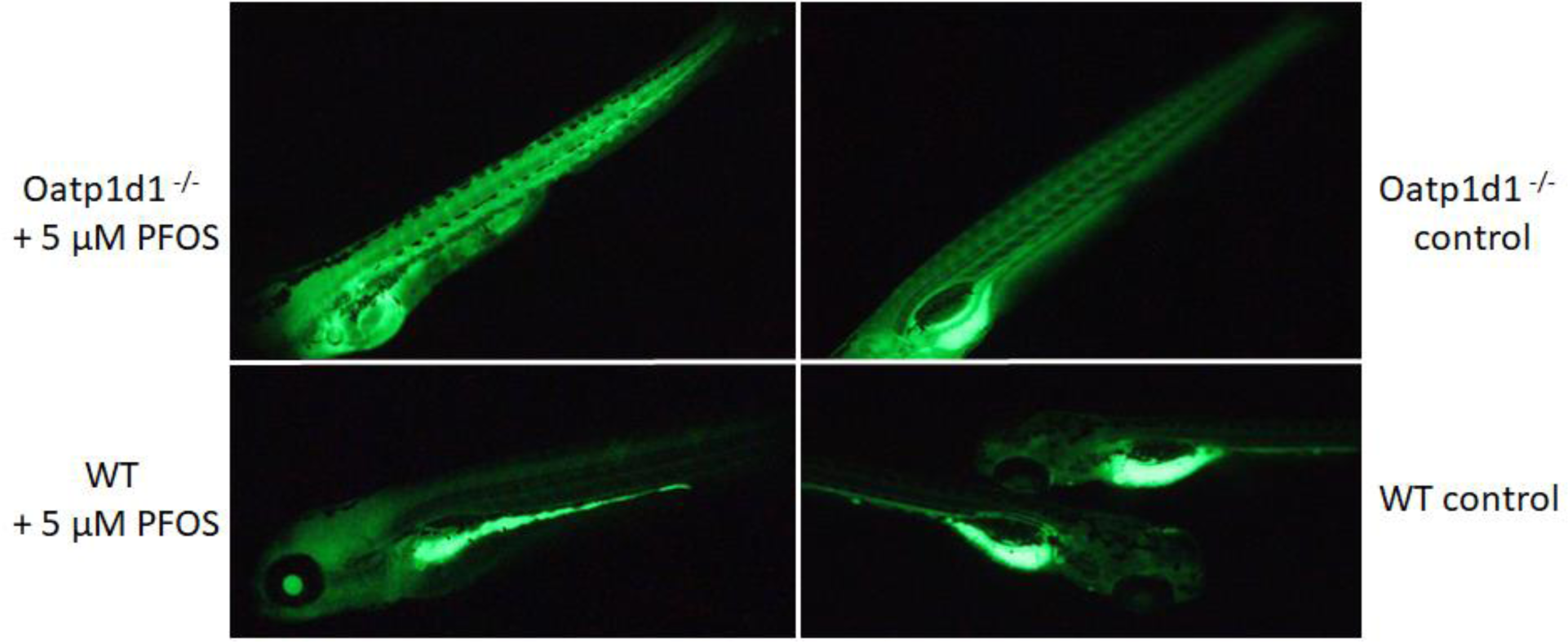
PFOS (5 µM) triggered oxidative stress during the initial stages of embryonal development. Representative images showing ROS production in WT and mutant embryos, stained with DCFH-DA.

**Figure 10.**
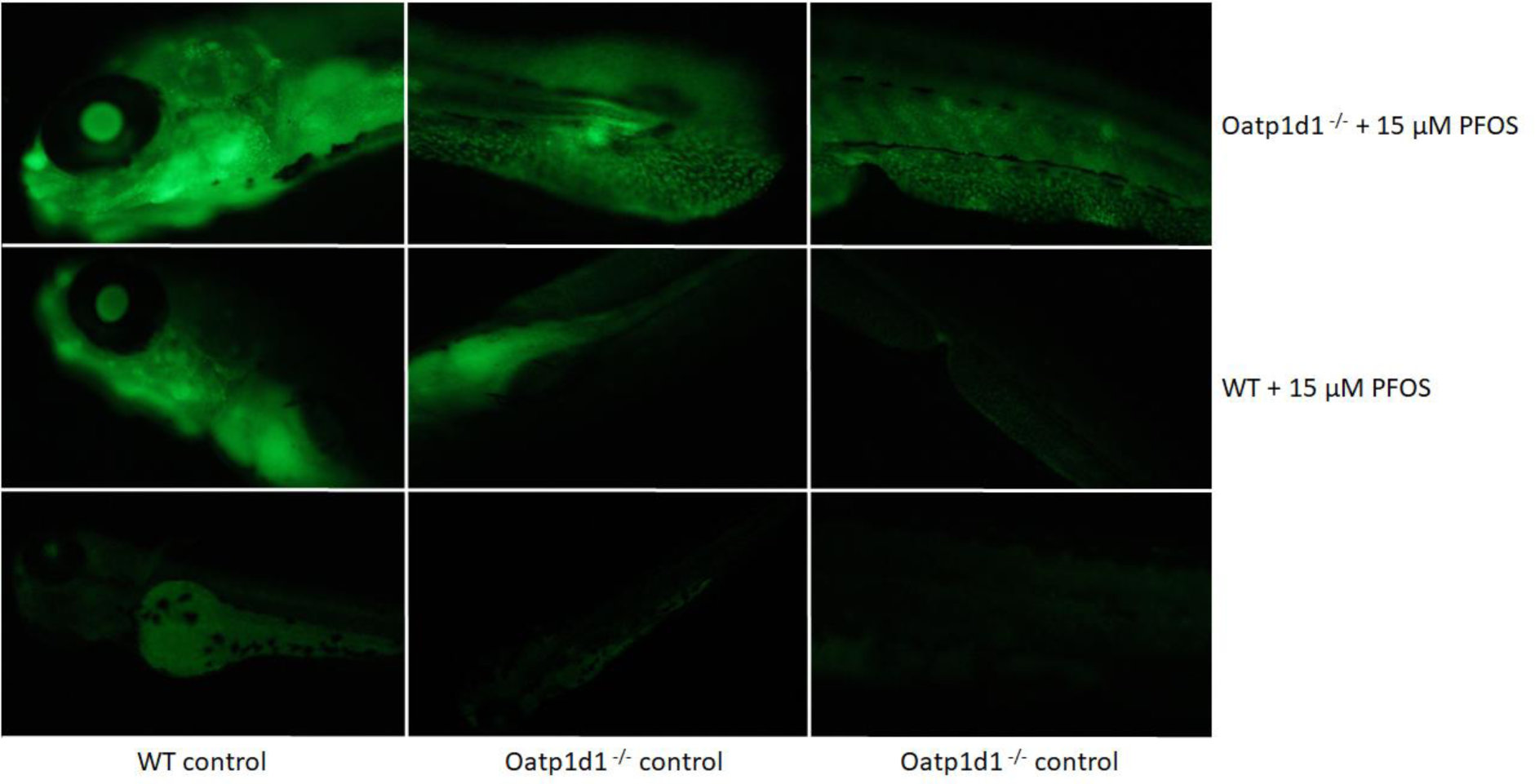
Apoptosis analysis following 15 µM PFOS exposure at 72 hpf. Representative images of apoptotic cells in the head and tail section, stained with acridine orange.

*Abcg2a* showed 1.5-fold upregulation in 5 dpf wt embryos after exposure to 15 µM PFOS for 1 h and a 1.14-fold upregulation in mutant embryos (Fig. 5). Longer exposure to 5 µM PFOS caused no changes in the transcript expression of abcg2a in both wt and mutant embryos (Fig. 6). However, at 4 dpf wt embryos showed high downregulation of 0.26-fold in wt and 0.46-fold in mutant embryos after three days of exposure to 30 µM PFOS (Fig. 7).

The *cyp1a* gene showed generally low expression under all conditions, with weak modulations of transcript expression upon exposure to different PFOS concentrations (Figs. 6, 8). *Cyp3a* showed a 2.16-fold upregulation in 5 dpf wt embryos after exposure to 15 µM PFOS for 1 h, whereas mutant embryos showed higher upregulation of 4.78-fold (Fig. S1). Longer exposure to 5 µM PFOS caused 0.75-fold downregulation of cyp3a transcript in wt embryos, and 3.12-fold upregulation in mutant embryos (Fig. S2). However, at 4 dpf, wt embryos showed high downregulation of 0.03-fold upon exposure to 30 µM PFOS for three days, and lower downregulation of 0.72 in mutant embryos (Fig. S3).

*Gstr1* gene showed generally low expression under all conditions, with weak modulations of transcript expression upon exposure to different concentrations of PFOS (Figs. S6). *Gstp1-2* genes showed downregulation after exposure with 15 µM PFOS for 1 h of 0.78-fold and 0.72-fold in wt and mutant embryos, respectively (Fig. S4). Longer exposure with 5 µM PFOS caused 0.84- fold downregulation of *gstp1-2* transcript in WT embryos and 0.77-fold downregulation in mutant embryos (Fig. S4). 4 dpf WT embryos showed downregulation of 0.81-fold upon exposure to 30 µM PFOS for three days, and no modulation of transcript expression in mutant embryos (Fig. S5).

## DISCUSSION

This study presents a comprehensive analysis of the impacts of perfluorooctane sulfonate (PFOS) exposure on zebrafish embryos, highlighting the significant differences in response between wild-type (WT) and *oatp1d1* mutant embryos. The findings contribute to our understanding of the toxicokinetic and toxicodynamic mechanisms of PFOS, a prominent member of the per- and polyfluoroalkyl substances (PFAS) family known for their environmental persistence and bioaccumulative potential.

The mortality rates observed in WT and *oatp1d1* mutant embryos following PFOS exposure reveal crucial involvement of Oatp1d1-mediated transport into the differential sensitivity of these two groups. The calculated LC_50_ values, 23.57 µM for WT and 16.71 µM for *oatp1d1* mutants, indicate that the mutants are more susceptible to PFOS toxicity. This difference underscores the critical role of the Oatp1d1 transporter in mediating the toxic effects of PFOS. The higher mortality rate in the mutant embryos suggests that Oatp1d1 may be involved in the detoxification processes or in the regulation of PFOS bioavailability, a finding consistent with previous research highlighting the importance of transporters in managing PFOS toxicity (Dragojević et al., 2021; Popovic et al., 2014).

The swim bladder, a vital organ for buoyancy and balance in fish, is crucial for normal aquatic life and its impairment can lead to severe physiological consequences (Mylroie et al., 2021). Exposure of developing zebrafish embryos to PFOS can lead to significant developmental abnormalities, with the most prominent effects observed in the swim bladder (Hagenaars et al., 2014). The absence or reduction in size of the swim bladder in exposed embryos resulted in reduced activity and abnormal positioning, e.g. when embryos lay on their side. These effects were more severe in *oatp1d1* mutant embryos, suggesting that the Oatp1d1 transporter plays a crucial role in defense against PFOS-induced developmental toxicity. The developmental abnormalities are consistent with previous studies showing that PFOS can disrupt normal organ development (S. Wang et al., 2017). The ability of PFOS to interfere with crucial developmental pathways can lead to a range of abnormalities, from impaired swim bladder development to scoliosis and necrosis in the head and gut, particularly in mutant embryos (Lau et al., 2007). These findings emphasize the need for further research into the specific mechanisms by which PFOS disrupts developmental processes.

This study provides a detailed analysis of the changes in gene expression induced by PFOS exposure, focusing on genes involved in biotransformation processes. The upregulation of *oatp1d1* in WT embryos after PFOS exposure contrasts with the lack of changes in mutant embryos, which has already been observed in the functional knockout of the Oatp1d1 transporter (Vujica et al., n.d.). This differential gene expression highlights the role of transporters in the biotransformation and detoxification of PFOS. Additionally, we observed significant modulations in the expression of *oatp2b1*, another important transporter from the same Slc21 family (Dragojević et al., 2021). In both WT and mutant embryos, PFOS exposure led to an upregulation of *oatp2b1*, although the response was slightly more pronounced in the mutant embryos. These findings suggest that Oatp2b1 may also play an important role in mediating the toxic effects of PFOS. The involvement of multiple transporters in PFOS toxicity underscores the complexity of its toxicokinetics and the need for a comprehensive understanding of these mechanisms in order to develop effective mitigation strategies.

PFOS exposure affects the expression of cytochrome P450 (*cyp*) and glutathione S-transferase (*gst*) genes. These genes are crucial for the detoxification processes and the metabolism of xenobiotics (Esteves et al., 2021). Cytochrome P450 enzymes encoded by the *cyp* family of genes play a pivotal role in the oxidative metabolism of a wide range of endogenous and exogenous compounds (Nawaji et al., 2020). We observed modulations in the expression of several *cyp* genes, indicating that PFOS exposure can alter the metabolic capacity of zebrafish embryos. The low expression of *cyp1a* with low inducibility was expected due to the higher expression of *cyp3a*, which is more involved in detoxification processes during early zebrafish development, whereas *cyp1a* involvement increases in adult zebrafish (Verbueken et al., 2017). The upregulation of *cyp3a* in WT embryos after PFOS exposure suggests an adaptive response to increase detoxification and metabolic processing of PFOS. However, the different expression patterns between WT and mutant embryos, with the mutants showing higher upregulation of *cyp3a*, highlight the potential compensatory mechanisms activated in response to PFOS-induced stress. These findings are consistent with previous studies that have documented the induction of cytochrome P450 enzymes in response to chemical stressors that may enhance the organism’s ability to metabolize and eliminate toxicants (Hahn E. & Stegeman J., 1994).

Similarly, glutathione S-transferase enzymes, encoded by *gst* genes, are involved in the conjugation of glutathione to a wide range of substrates and facilitate their excretion (Glisic et al., 2015). In our study, changes in the expression of the *gstp1-2* and *gstr1* genes were observed upon PFOS exposure. The downregulation of these genes in both WT and mutant embryos suggests a compromised ability to detoxify reactive intermediates generated by PFOS metabolism. This downregulation could lead to increased oxidative stress and cellular damage, as glutathione conjugation is a critical mechanism for neutralizing reactive oxygen species and preventing oxidative damage (Hayes & Pulford, 1995). This is something that we observed in this study, as *oatp1d1* mutant embryos showed higher levels of oxidative stress compared to WT embryos. Additionally, we observed higher levels of apoptosis in mutant embryos, suggesting that the loss of Oatp1d1might lead to inefficient excretion of PFOS and toxic levels in the bloodstream that could cause deleterious developmental malformations.

## CONCLUSION

The observed gene expression changes, oxidative stress and apoptosis levels emphasize the complexity of PFOS toxicity and highlight the versatile nature of the organism’s response to PFOS-induced stress. The differential modulation of membrane transporter genes as well as *cyp* and *gst* genes between WT and mutant embryos provides valuable insights into the specific pathways affected by PFOS and the potential genetic factors that influence susceptibility to PFOS toxicity. This study provides a detailed examination of the toxic effects of PFOS on zebrafish embryos highlighting the critical role of transporters in mediating these effects. The differential sensitivity of WT and *oatp1d1* mutant embryos to PFOS exposure underscores the importance of genetic factors in determining toxicity. The findings from this study have significant implications for aquatic ecosystems, particularly regarding the environmental persistence and bioaccumulative nature of PFOS. The observed developmental and physiological abnormalities in zebrafish embryos highlight the potential risks to aquatic organisms, particularly at higher trophic levels. The bioaccumulation of PFOS in aquatic organisms may lead to higher concentrations in predators, posing risks to wildlife and potentially impacting entire aquatic food webs. The study’s insights into the molecular mechanisms of PFOS toxicity provide a basis for understanding the broader ecological impacts of PFOS contamination. The differential sensitivity of WT and oatp1d1 mutant embryos to PFOS exposure underscores the importance of considering genetic and species-specific differences when assessing environmental risks.

## Authorship contribution statement

I. Mihaljevic: Conceptualization, data curation, formal analysis, investigation, methodology, visualization, validation, writing – original draft. L. Vujica: Conceptualization, data curation, formal analysis, investigation, methodology, writing – original draft. J. Dragojevic: data curation, formal analysis, investigation, methodology, writing – original draft. J. Loncar: Data curation, formal analysis, investigation, methodology, writing – original draft. T. Smital: Conceptualization, writing – review & editing, validation, supervision, funding acquisition.

## Supplementary Material

**Figure S1.**
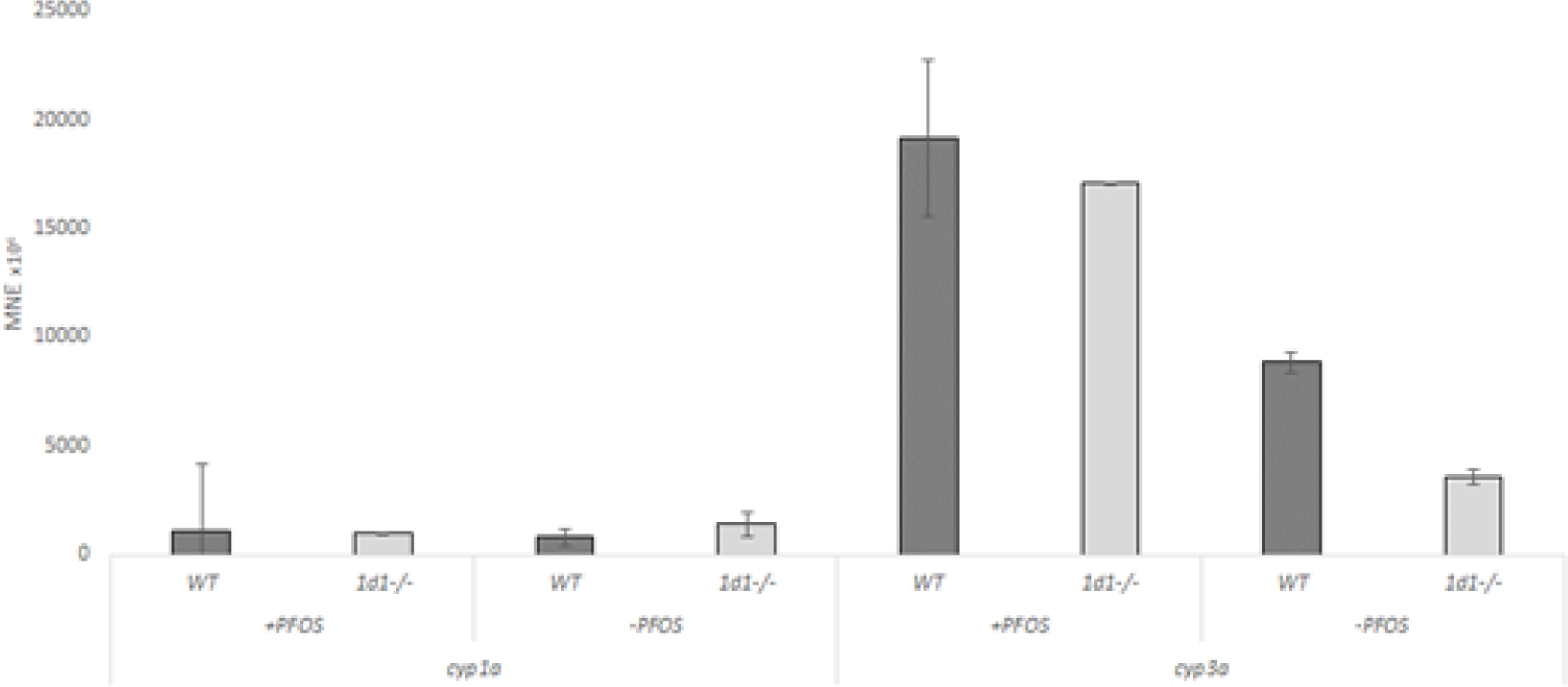
Transcript expression patterns of *cyp1a* and *cyp3a* in WT and *oatp1d1* mutant embryos at 5 dpf and with and without 1h exposure to 15 µM PFOS. Results of pools of 10 embryos per sample are presented. Data represents MNE (mean normalized expression) ± SD normalized to the housekeeping gene *ATP50*.

**Figure S2.**
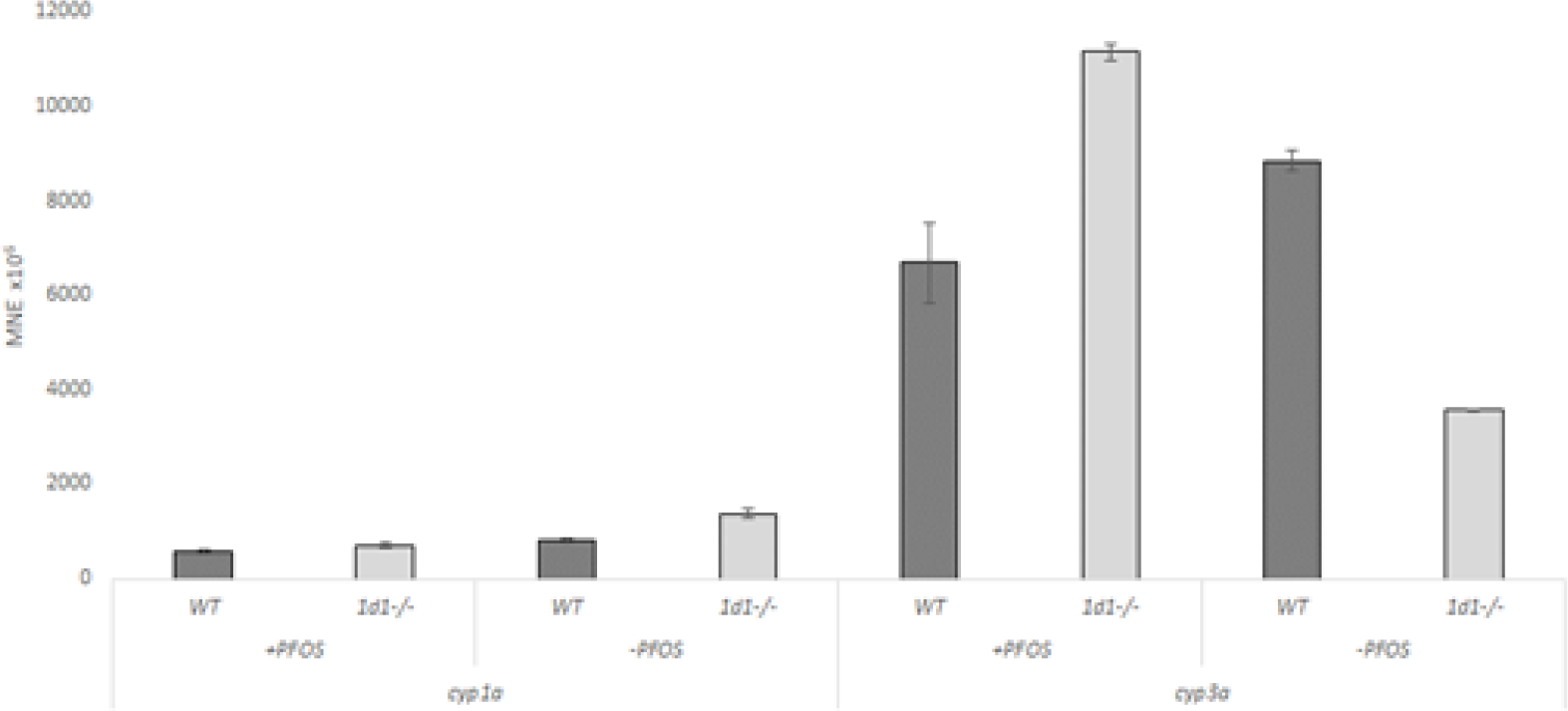
Transcript expression patterns of *cyp1a* and *cyp3a* in WT and *oatp1d1* mutant embryos at 5 dpf and with and without 4 days exposure to 5 µM PFOS. Results of pools of 10 embryos per sample are presented. Data represents MNE (mean normalized expression) ± SD normalized to the housekeeping gene *ATP50*.

**Figure S3.**
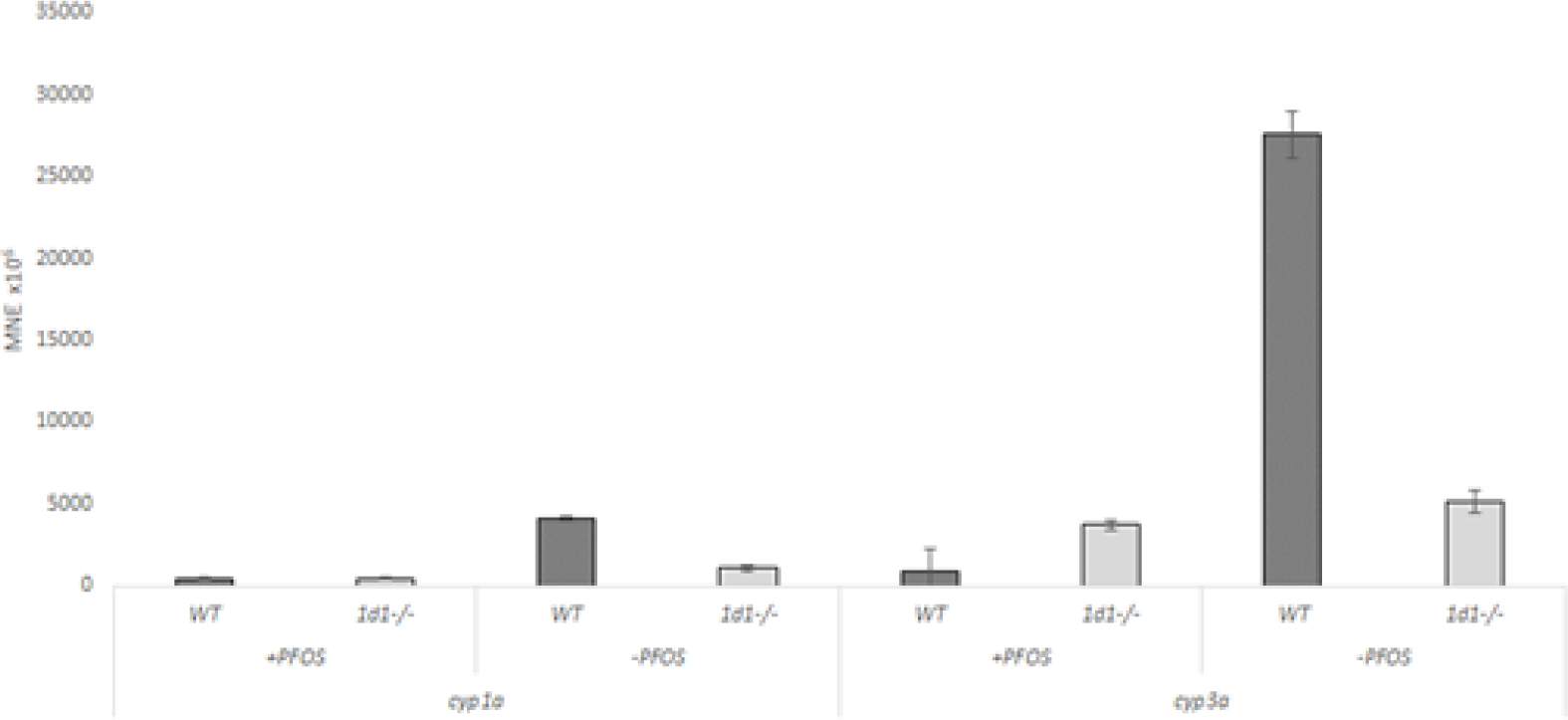
Transcript expression patterns of *cyp1a* and *cyp3a* in WT and *oatp1d1* mutant embryos at 4 dpf and with and without 3 days exposure to 30 µM PFOS. Results of pools of 10 embryos per sample are presented. Data represents MNE (mean normalized expression) ± SD normalized to the housekeeping gene *ATP50*.

**Figure S4.**
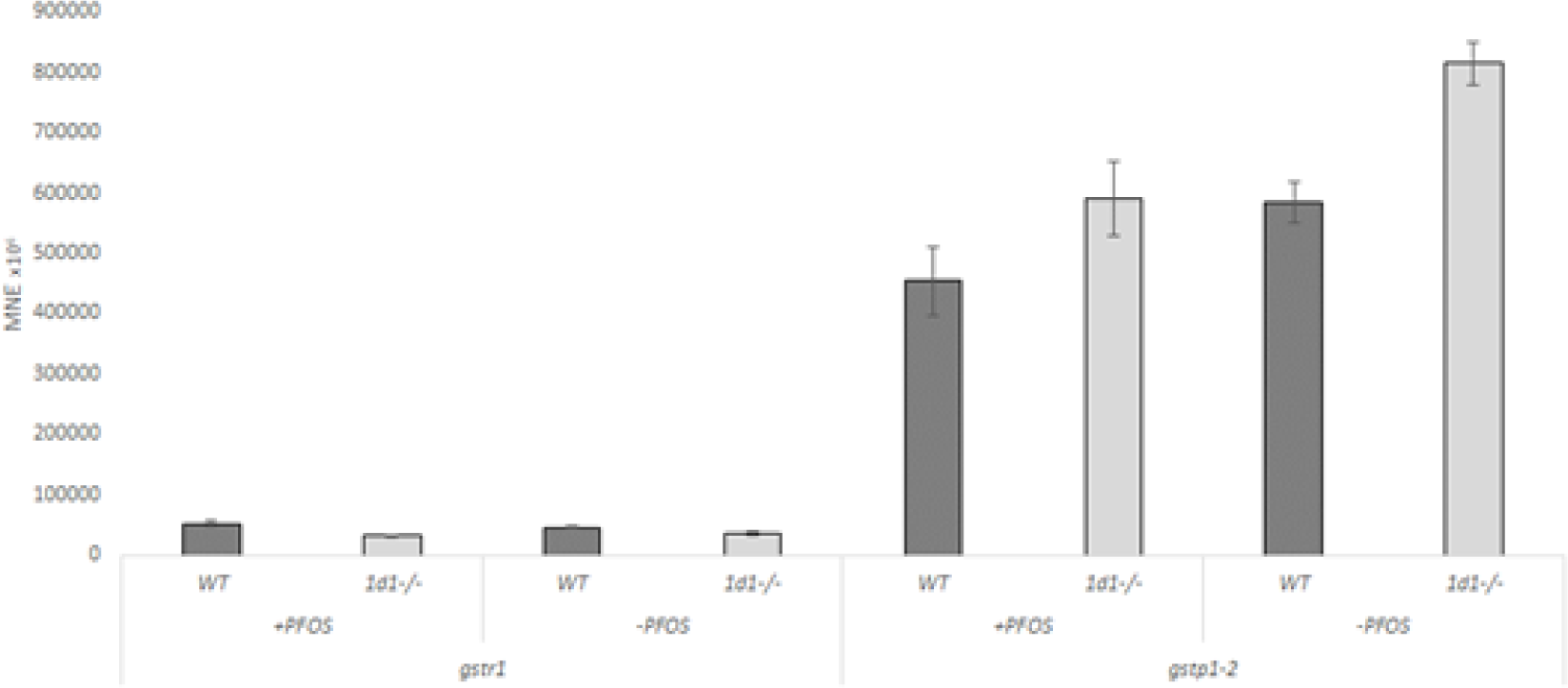
Transcript expression patterns of *gstr1* and *gstp1-2* in WT and *oatp1d1* mutant embryos at 5 dpf and with and without 1 h exposure to 15 µM PFOS. Results of pools of 10 embryos per sample are presented. Data represents MNE (mean normalized expression) ± SD normalized to the housekeeping gene *ATP50*.

**Figure S5.**
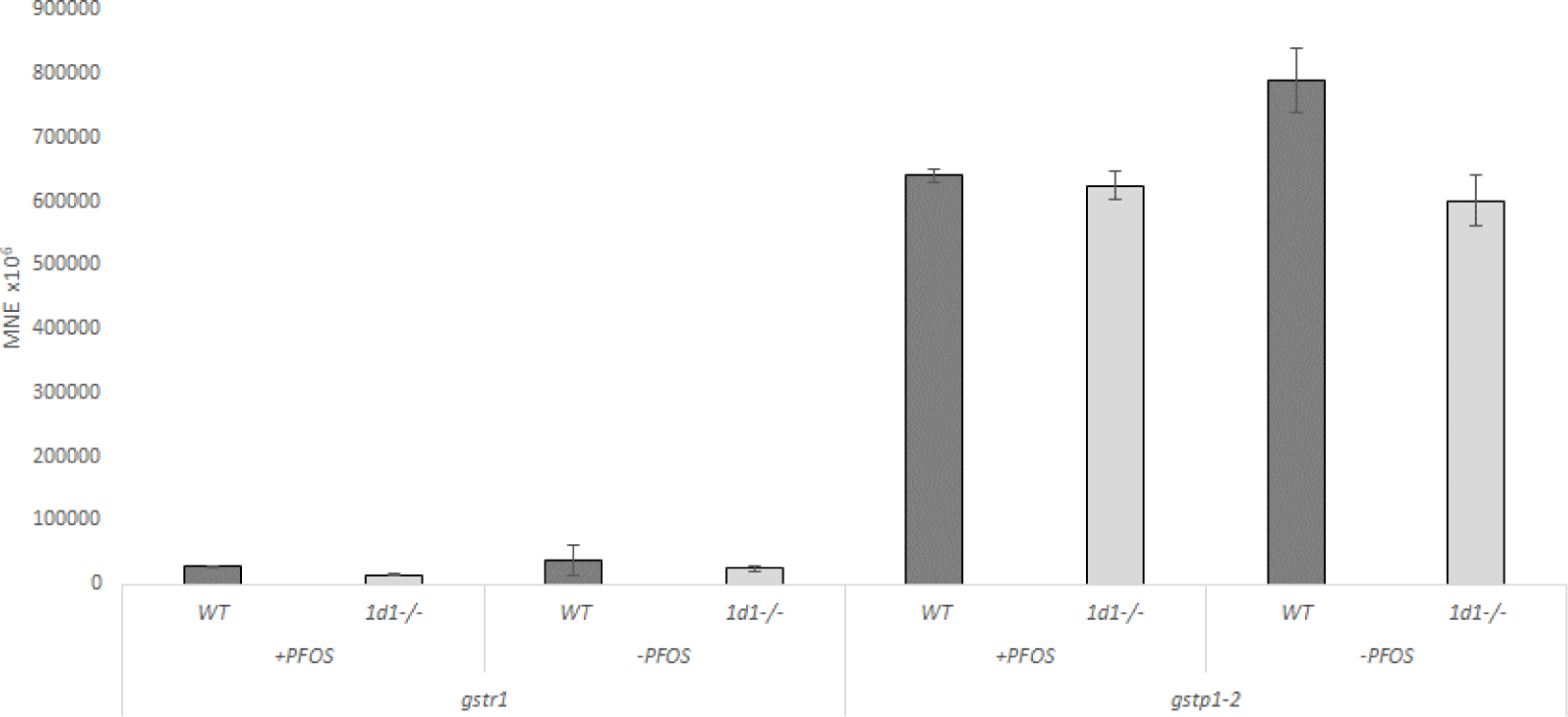
Transcript expression patterns of *gstr1* and *gstp1-2* in WT and *oatp1d1* mutant embryos at 4 dpf and with and without 3 days exposure to 30 µM PFOS. Results of pools of 10 embryos per sample are presented. Data represents MNE (mean normalized expression) ± SD normalized to the housekeeping gene *ATP50*.

**Figure S6.**
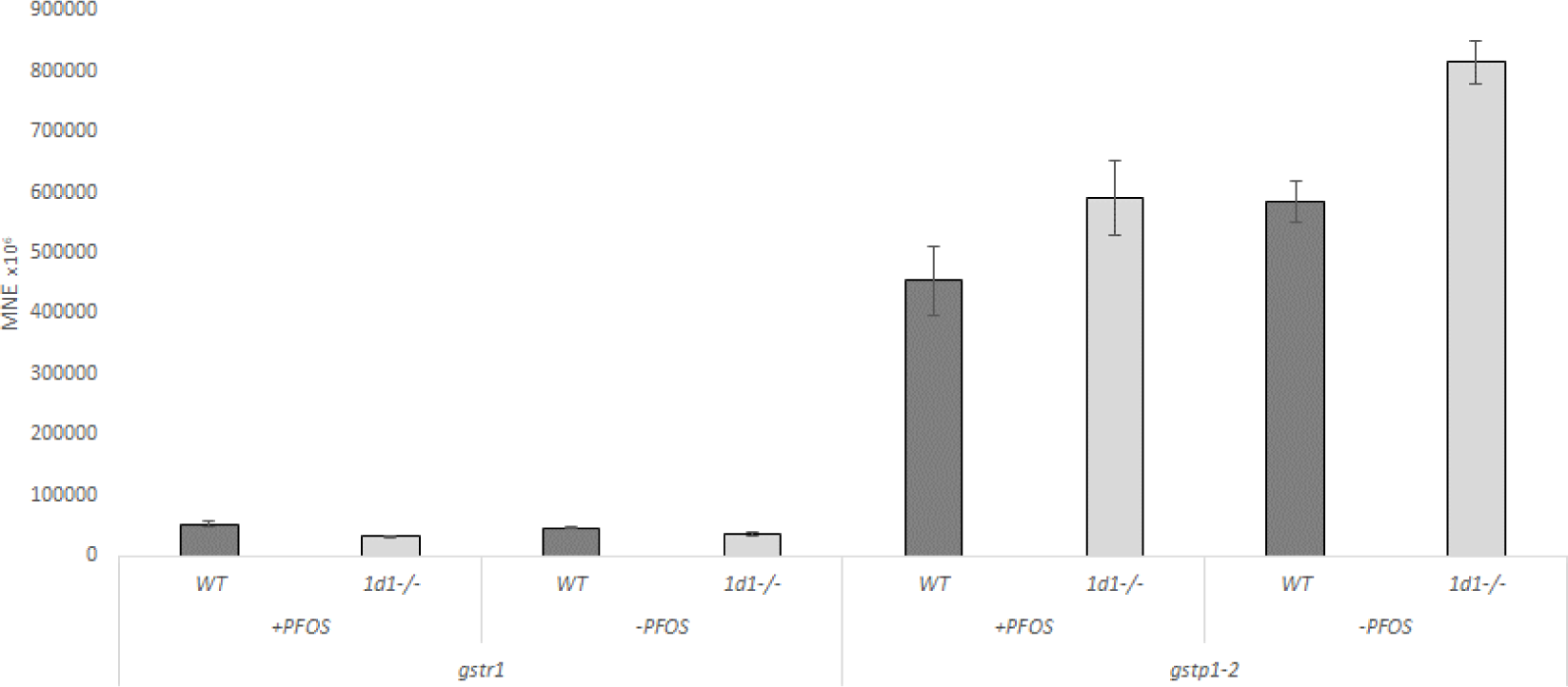
Transcript expression patterns of *gstr1* and *gstp1-2* in WT and *oatp1d1* mutant embryos at 5 dpf and with and without 1 h exposure to 15 µM PFOS. Results of pools of 10 embryos per sample are presented. Data represents MNE (mean normalized expression) ± SD normalized to the housekeeping gene *ATP50*.

**Figure S7.**
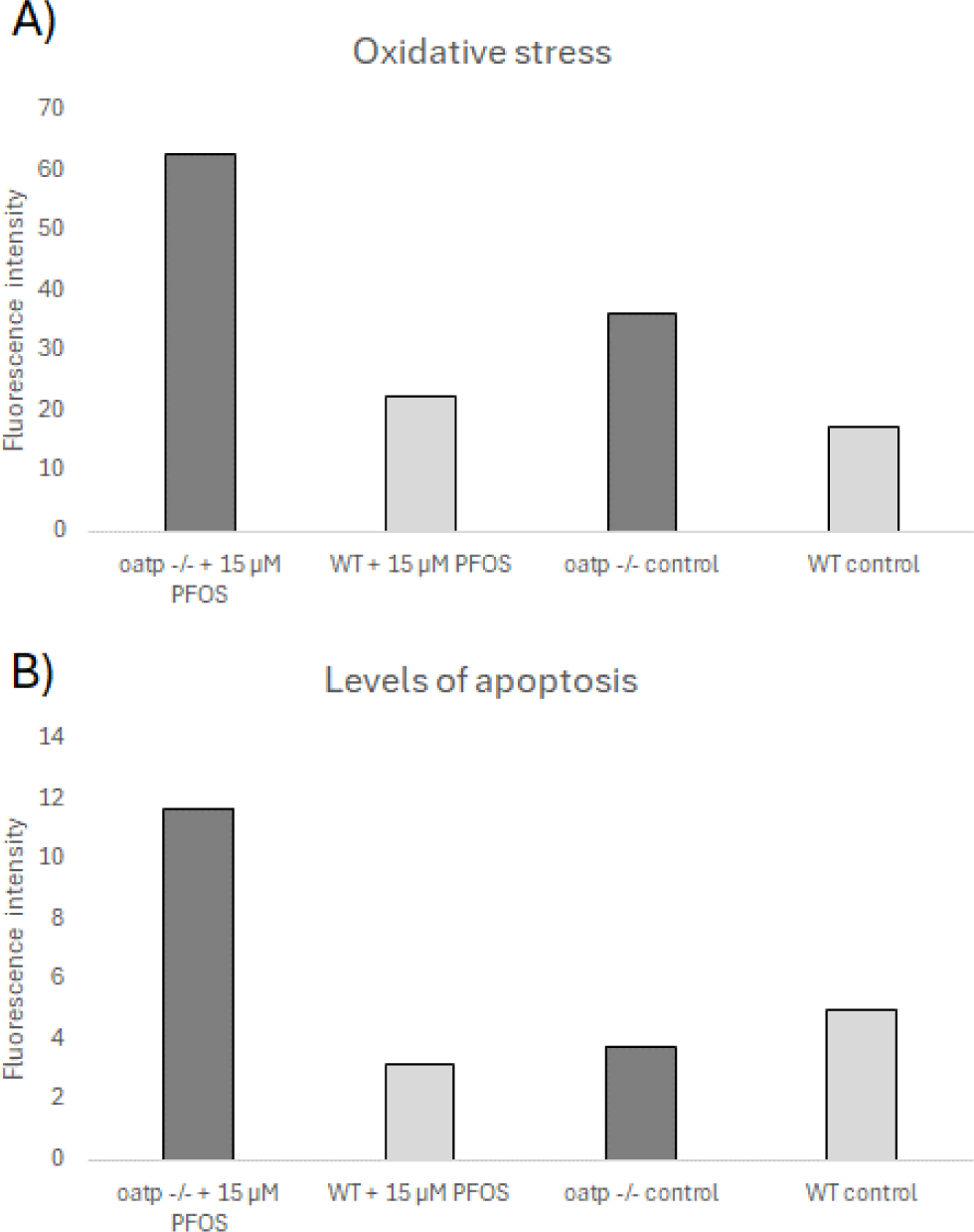
Levels of oxidative stress and apoptosis following PFOS exposure of both WT and *oatp1d1* mutant embryos. (A) ROS levels as fluorescence intensity obtained by H_2_DCFDA method showing higher levels of oxidative stress in mutant embryos treated with 15 µM PFOS. (B) Apoptosis levels obtained by acridine organge method showing simila pattern and higher levels in mutant embryos exposed to 15 µM PFOS.

**Figure S8.**
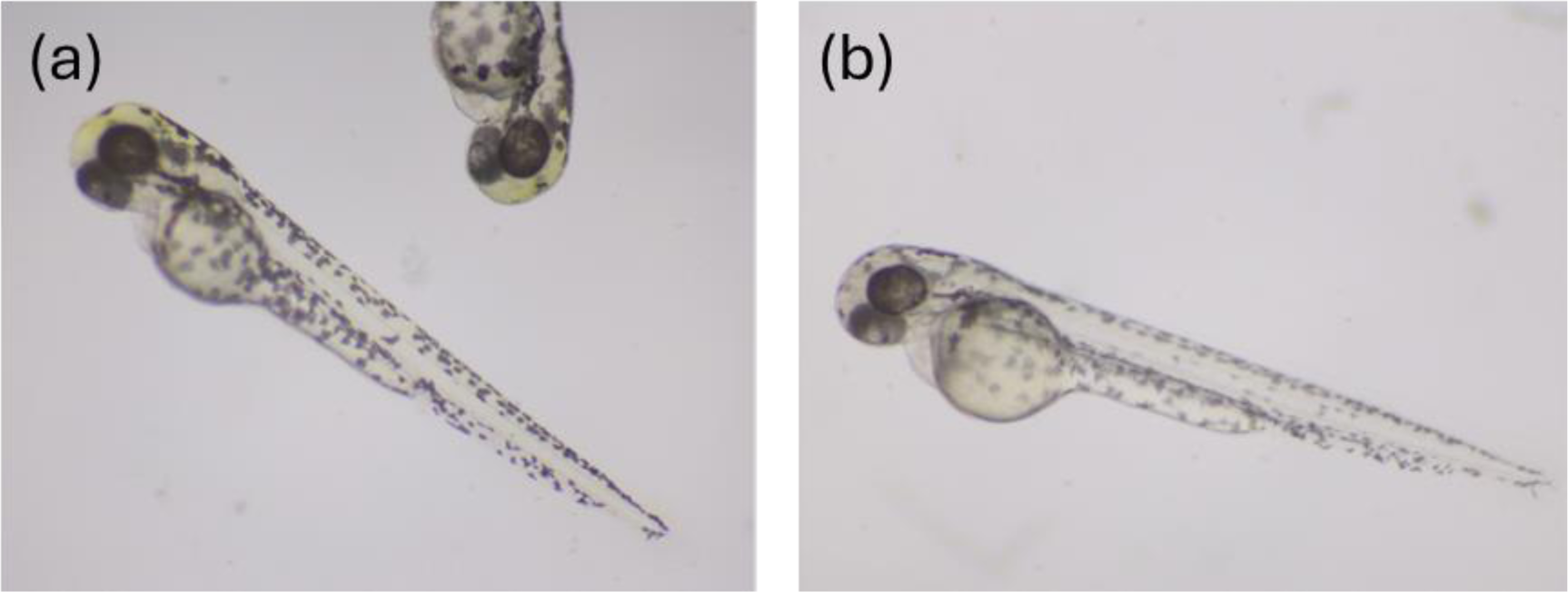
Images of 4 dpf DMSO treated control embryos. (a) wild type, (b) *oatp1d1* mutant.

## REFERENCES

Ahrens, L. (2011). Polyfluoroalkyl compounds in the aquatic environment: A review of their occurrence and fate. Journal of Environmental Monitoring, 13(1), 20–31. 10.1039/c0em00373e

Ankley, G. T., Cureton, P., Hoke, R. A., Houde, M., Kumar, A., Kurias, J., Lanno, R., McCarthy, C., Newsted, J., Salice, C. J., Sample, B. E., Sepúlveda, M. S., Steevens, J., & Valsecchi, S. (2021). Assessing the Ecological Risks of Per- and Polyfluoroalkyl Substances: Current State-of-the Science and a Proposed Path Forward. In Environmental Toxicology and Chemistry (Vol. 40, Issue 3, pp. 564–605). Wiley Blackwell. 10.1002/etc.4869

Buck, R. C., Franklin, J., Berger, U., Conder, J. M., Cousins, I. T., Voogt, P. De, Jensen, A. A., Kannan, K., Mabury, S. A., & van Leeuwen, S. P. J. (2011). Perfluoroalkyl and polyfluoroalkyl substances in the environment: Terminology, classification, and origins. Integrated Environmental Assessment and Management, 7(4), 513–541. 10.1002/ieam.258

Conder, J. M., Hoke, R. A., De Wolf, W., Russell, M. H., & Buck, R. C. (2008). Are PFCAs bioaccumulative? A critical review and comparison with regulatory criteria and persistent lipophilic compounds. In Environmental Science and Technology (Vol. 42, Issue 4, pp. 995– 1003). 10.1021/es070895g

Consoer, D. M., Hoffman, A. D., Fitzsimmons, P. N., Kosian, P. A., & Nichols, J. W. (2016). Toxicokinetics of perfluorooctane sulfonate in rainbow trout (Oncorhynchus mykiss). Environmental Toxicology and Chemistry, 35(3), 717–727. 10.1002/etc.3230

Cousins, I. T., Vestergren, R., Wang, Z., Scheringer, M., & McLachlan, M. S. (2016). The precautionary principle and chemicals management: The example of perfluoroalkyl acids in groundwater. In Environment International (Vol. 94, pp. 331–340). Elsevier Ltd. 10.1016/j.envint.2016.04.044

Domingo, J. L., & Nadal, M. (2019). Human exposure to per- and polyfluoroalkyl substances (PFAS) through drinking water: A review of the recent scientific literature. In Environmental Research (Vol. 177). Academic Press Inc. 10.1016/j.envres.2019.108648

Dragojević, J., Marakovic, N., Popović, M., & Smital, T. (2021). Zebrafish (Danio rerio) Oatp2b1 as a functional ortholog of the human OATP2B1 transporter. Fish Physiology and Biochemistry, 47(6), 1837–1849. 10.1007/s10695-021-01015-7

Dragojević, J., Marić, P., Lončar, J., Popović, M., Mihaljević, I., & Smital, T. (2020). Environmental contaminants modulate transport activity of zebrafish organic anion transporters Oat1 and Oat3. Comparative Biochemistry and Physiology Part - C: Toxicology and Pharmacology, 231. 10.1016/j.cbpc.2020.108742

Esteves, F., Rueff, J., & Kranendonk, M. (2021). The central role of cytochrome p450 in xenobiotic metabolism—a brief review on a fascinating enzyme family. In Journal of Xenobiotics (Vol. 11, Issue 3, pp. 94–114). MDPI. 10.3390/jox11030007

Giesy, J. P., & Kannan, K. (2001). Global distribution of perfluorooctane sulfonate in wildlife. Environmental Science and Technology, 35(7), 1339–1342. 10.1021/es001834k

Glisic, B., Mihaljevic, I., Popovic, M., Zaja, R., Loncar, J., Fent, K., Kovacevic, R., & Smital, T. (2015). Characterization of glutathione-S-transferases in zebrafish (Danio rerio). Aquatic Toxicology, 158, 50–62. 10.1016/j.aquatox.2014.10.013

Hagenaars, A., Stinckens, E., Vergauwen, L., Bervoets, L., & Knapen, D. (2014). PFOS affects posterior swim bladder chamber inflation and swimming performance of zebrafish larvae. Aquatic Toxicology, 157, 225–235. 10.1016/j.aquatox.2014.10.017

Hahn E., M., & Stegeman J., J. (1994). Regulation of Cytochrome P4501A1 in Teleosts: Sustained Induction of CYP1A1 mRNA, Protein, and Catalytic Activity by 2,3,7,8-Tetrachlorodibenzofuran in the Marine Fish Stenotomus chrysops. Toxicology and Applied Pharmacology, 127, 187–198.

Hayes, J. D., & Pulford, D. J. (1995). The Glutathione S-Transferase Supergene Family: Regulation of GST* and the Contribution of the lsoenzymes to Cancer Chemoprotection and Drug Resistance. In Critical Reviews in Biochemistry and Molecular Biology Downloaded from informahealthcare (Vol. 30, Issue 6).

Houde, M., Martin, J. W., Letcher, R. J., Solomon, K. R., & Muir, D. C. G. (2006). Biological monitoring of polyfluoroalkyl substances: A review. In Environmental Science and Technology (Vol. 40, Issue 11, pp. 3463–3473). 10.1021/es052580b

Jantzen, C. E., Annunziato, K. M., & Cooper, K. R. (2016). Behavioral, morphometric, and gene expression effects in adult zebrafish (Danio rerio) embryonically exposed to PFOA, PFOS, and PFNA. Aquatic Toxicology, 180, 123–130. 10.1016/j.aquatox.2016.09.011

Kimmel, C. B., Ballard, W. W., Kimmel, S. R., Ullmann, B., & Schilling, T. F. (1995). Stages of embryonic development of the zebrafish. Developmental Dynamics, 203(3), 253–310. 10.1002/aja.1002030302

Kimura, O., Fujii, Y., Haraguchi, K., Kato, Y., Ohta, C., Koga, N., & Endo, T. (2020). Effects of perfluoroalkyl carboxylic acids on the uptake of sulfobromophthalein via organic anion transporting polypeptides in human intestinal Caco-2 cells. Biochemistry and Biophysics Reports, 24. 10.1016/j.bbrep.2020.100807

Lau, C., Anitole, K., Hodes, C., Lai, D., Pfahles-Hutchens, A., & Seed, J. (2007). Perfluoroalkyl acids: A review of monitoring and toxicological findings. In Toxicological Sciences (Vol. 99, Issue 2, pp. 366–394). 10.1093/toxsci/kfm128

Lončar, J., Popović, M., Zaja, R., & Smital, T. (2010). Gene expression analysis of the ABC efflux transporters in rainbow trout (Oncorhynchus mykiss). Comparative Biochemistry and Physiology - C Toxicology and Pharmacology, 151(2), 209–215. 10.1016/j.cbpc.2009.10.009

Lou, Q. Q., Zhang, Y. F., Zhou, Z., Shi, Y. L., Ge, Y. N., Ren, D. K., Xu, H. M., Zhao, Y. X., Wei, W. J., & Qin, Z. F. (2013). Effects of perfluorooctanesulfonate and perfluorobutanesulfonate on the growth and sexual development of Xenopus laevis. Ecotoxicology, 22(7), 1133–1144. 10.1007/s10646-013-1100-y

Mihaljevic, I., Popovic, M., Zaja, R., & Smital, T. (2016). Phylogenetic, syntenic, and tissue expression analysis of slc22 genes in zebrafish (Danio rerio). BMC Genomics, 17(1). 10.1186/s12864-016-2981-y

Modzelewski, A. J., Chen, S., Willis, B. J., Lloyd, K. C. K., Wood, J. A., & He, L. (2018). Efficient mouse genome engineering by CRISPR-EZ technology. Nature Protocols, 13(6), 1253–1274. 10.1038/nprot.2018.012

Mylroie, J. E., Wilbanks, M. S., Kimble, A. N., To, K. T., Cox, C. S., McLeod, S. J., Gust, K. A., Moore, D. W., Perkins, E. J., & Garcia-Reyero, N. (2021). Perfluorooctanesulfonic Acid– Induced Toxicity on Zebrafish Embryos in the Presence or Absence of the Chorion. Environmental Toxicology and Chemistry, 40(3), 780–791. 10.1002/etc.4899

Nakagawa, H., Terada, T., Harada, K. H., Hitomi, T., Inoue, K., Inui, K. I., & Koizumi, A. (2009). Human organic anion transporter hOAT4 is a transporter of perfluorooctanoic acid. Basic and Clinical Pharmacology and Toxicology, 105(2), 136–138. 10.1111/j.1742-7843.2009.00409.x

Nawaji, T., Yamashita, N., Umeda, H., Zhang, S., Mizoguchi, N., Seki, M., Kitazawa, T., & Teraoka, H. (2020). Cytochrome P450 expression and chemical metabolic activity before full liver development in Zebrafish. Pharmaceuticals, 13(12), 1–17. 10.3390/ph13120456

Popovic, M., Zaja, R., Fent, K., & Smital, T. (2013). Molecular characterization of zebrafish Oatp1d1 (Slco1d1), a novel organic anion-transporting polypeptide. Journal of Biological Chemistry, 288(47), 33894–33911. 10.1074/jbc.M113.518506

Popovic, M., Zaja, R., Fent, K., & Smital, T. (2014). Interaction of environmental contaminants with zebrafish organic anion transporting polypeptide, Oatp1d1 (Slco1d1). *Toxicology and Applied Pharmacology*, *280*(1), 149–158. 10.1016/j.taap.2014.07.015

Post, G. B., Cohn, P. D., & Cooper, K. R. (2012). Perfluorooctanoic acid (PFOA), an emerging drinking water contaminant: A critical review of recent literature. In Environmental Research (Vol. 116, pp. 93–117). 10.1016/j.envres.2012.03.007

Prevedouros, K., Cousins, I. T., Buck, R. C., & Korzeniowski, S. H. (2006). Sources, fate and transport of perfluorocarboxylates. In Environmental Science and Technology (Vol. 40, Issue 1, pp. 32–44). 10.1021/es0512475

Rosenmai, A. K., Taxvig, C., Svingen, T., Trier, X., van Vugt-Lussenburg, B. M. A., Pedersen, M., Lesné, L., Jégou, B., & Vinggaard, A. M. (2016). Fluorinated alkyl substances and technical mixtures used in food paper-packaging exhibit endocrine-related activity in vitro. Andrology, 4(4), 662–672. 10.1111/andr.12190

Sato, I., Kawamoto, K., Nishikawa, Y., Shuji, T., Yoshida, M., Yaegashi, K., Saito, N., Liu, W., & Jin, Y. (2009). Neurotoxicity of perflourooctane sulfonate (PFOS) in rats and mice after single oral exposure. The Journal of Toxicological Sciences, 34(5), 569–574.

Sun, M., Arevalo, E., Strynar, M., Lindstrom, A., Richardson, M., Kearns, B., Pickett, A., Smith, C., & Knappe, D. R. U. (2016). Legacy and Emerging Perfluoroalkyl Substances Are Important Drinking Water Contaminants in the Cape Fear River Watershed of North Carolina. Environmental Science and Technology Letters, 3(12), 415–419. 10.1021/acs.estlett.6b00398

Verbueken, E., Alsop, D., Saad, M. A., Pype, C., van Peer, E. M., Casteleyn, C. R., Van Ginneken, C. J., Wilson, J., & Van Cruchten, S. J. (2017). In vitro biotransformation of two human CYP3A probe substrates and their inhibition during early zebrafish development. International Journal of Molecular Sciences, 18(1). 10.3390/ijms18010217

Vujica, L., Mihaljević, I., Dragojević, J., Lončar, J., Karaica, D., Bošnjak, A., & Smital, T. (n.d.). Functional knockout of the Oatp1d1 membrane transporter affects toxicity of 2 diclofenac in zebrafish embryos. https://ssrn.com/abstract=4811606

Wang, S., Zhuang, C., Du, J., Wu, C., & You, H. (2017). The presence of MWCNTs reduces developmental toxicity of PFOS in early life stage of zebrafish. Environmental Pollution, 222, 201–209. 10.1016/j.envpol.2016.12.055

Wang, Y. Q., Hu, L. X., Liu, T., Zhao, J. H., Yang, Y. Y., Liu, Y. S., & Ying, G. G. (2022). Per- and polyfluoralkyl substances (PFAS) in drinking water system: Target and non-target screening and removal assessment. Environment International, 163. 10.1016/j.envint.2022.107219

